# Neural representation of hedonic valence during narrative listening

**DOI:** 10.64898/2026.06.30.735538

**Authors:** Xuan Yang, Christian O’Reilly, Svetlana V. Shinkareva

**Affiliations:** Department of Psychology, University of South Carolina; Institute for Mind and Brain, University of South Carolina; Department of Computer Science and Engineering, University of South Carolina; Artificial Intelligence Institute, University of South Carolina; Carolina Autism and Neurodevelopment Research Center, University of South Carolina

**Keywords:** Valence, Naturalistic, Narrative listening, fMRI, Model Selection

## Abstract

Hedonic valence, the intrinsic pleasantness or unpleasantness of an experience, is fundamental to human psychological functioning. Yet, how valence is represented in the brain remains an open question. Functional MRI studies have demonstrated that the brain encodes both positive and negative valence, but this evidence largely stems from experiments using simplified, controlled stimuli, such as images, sounds, or words. As a result, it remains unclear how valence is processed during rich, naturalistic experiences that more closely reflect real life. In addition, most studies adopt a single statistical model, raising concerns about the robustness of their findings. This study used a formal voxel-wise Bayesian model selection approach to test alternative statistical models supporting Bipolarity, Valence-General, and Bivalence hypotheses to identify the most optimal model of valence representation during narrative listening. Our results provide evidence for the Bipolar model. We identified distributed brain regions that selectively encode valence as a bipolar continuum (negative to positive) during narrative comprehension, including classical emotion-related hubs such as ventromedial prefrontal cortex, as well as regions not traditionally associated with emotion processing, such as inferior occipital cortex, supramarginal cortex, inferior frontal cortex, and middle cingulate. Regions selectively encoding arousal and those broadly responsive to both valence and arousal were also identified. These findings highlight the importance of using formal model comparison and naturalistic paradigms in affective neuroscience, advancing our understanding of how valence is represented in the brain during real-world experiences.

---

Hedonic valence refers to the affective quality of pleasure or displeasure. It modulates nearly every aspect of psychological functioning, including perception, memory, judgment, language, social interaction, and well-being. Empirical studies using exploratory factor analysis and multidimensional scaling consistently identify valence as the most fundamental dimension of core affect and discrete emotions (Russell, 1980; Barrett & Bliss-Moreau, 2009). A deeper understanding of its neural representation is essential for advancing research in emotion, cognition, and mental health.

Valence is typically conceptualized as a continuum ranging from negative to positive affect. However, a long-standing debate persists over how valence is represented in the brain, whether positive and negative affect are encoded by the same or separate neural systems. Two primary theoretical accounts have been proposed (**Figure 1A**). The Bipolarity account posits that valence is represented along a single continuum from negative to positive affect (Russell, 2003; Russell & Barrett, 1999), with neural activity monotonically increasing or decreasing along this axis. Empirical support includes linear modulation of BOLD signals by valence in regions such as the orbitofrontal cortex (OFC), inferior frontal gyrus (IFG), superior and middle temporal gyri (STG, MTG), and insula in response to affective stimuli (Altmann et al., 2012; Anderson et al., 2003; Cunningham et al., 2004; Heinzel et al., 2005; Tseng et al., 2016).

**Figure 1.**
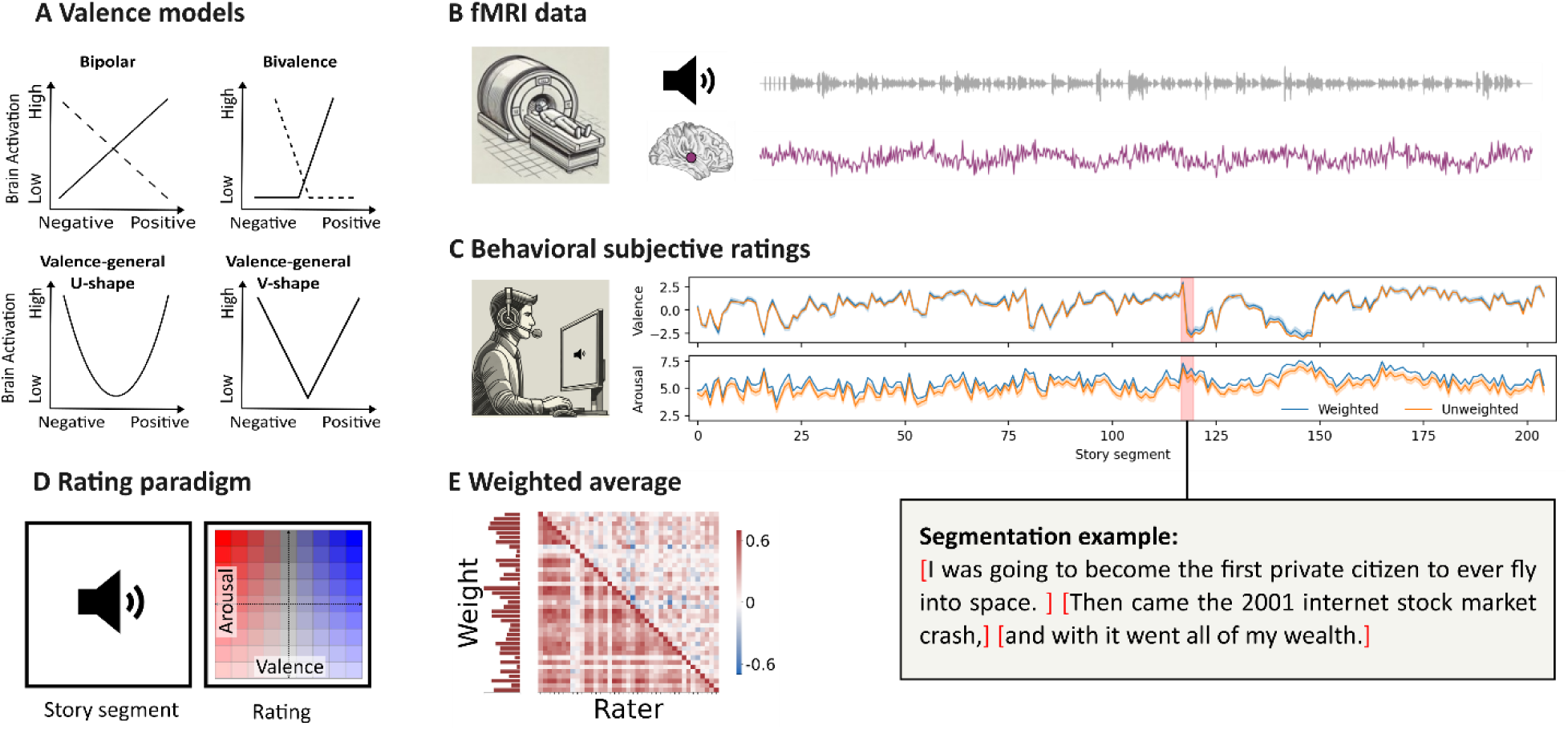
**A.** Hypothesized relationship between valence and brain activation for the four models. **B.** Illustration of the BOLD signal time course for a representative voxel during continuous narrative listening in the participants from the *Narratives* dataset. **C.** Illustration of subjective valence and arousal ratings from a separate sample of participants who listened to and rated the same narratives, one segment at a time. The line graphs show how valence and arousal evolve as the narrative unfolds. Blue and orange lines represent the weighted and unweighted average ratings, respectively, with the shaded area around the unweighted line indicating the 95% confidence interval. The highlighted portion of ratings corresponds to the narrative segments displayed in the bottom panel. **D.** Illustration of the subjective rating procedure for a single narrative segment. Participants first listened to a segment and then rated their affective response by clicking within a valence-by-arousal grid. **E.** Visualization of participant weights and intersubject rating similarity. *Left:* Bar graph of participant weights derived from the first eigenvector of RV-PCA applied to valence and arousal ratings. *Right:* Heatmap of pairwise Pearson correlations for a representative narrative (*Forgot*), with valence correlations shown in the lower triangle and arousal correlations in the upper triangle.

In contrast, the Bivalence account argues that although the behavioral manifestation of valence is bipolar, the underlying evaluation may not adhere to a bipolar structure (Cacioppo et al., 1997, 1999; Cacioppo & Berntson, 1994). Instead, positive and negative affect are thought to arise from distinct evaluative processes, encoded by independent neural systems. Although some studies report dissociable brain regions for each polarity, findings are inconsistent, with overlapping or divergent regions observed across tasks (Chikazoe et al., 2014; Lewis et al., 2006; Viinikainen et al., 2010). These inconsistencies raise questions about whether apparent selectivity reflects true neural specialization or limitations of classical statistical inference, as the absence of statistical significance does not confirm the absence of an effect.

In light of these mixed findings, some studies have instead reported a valence-general response profile in which neural activity increases with the magnitude of valence (i.e., equivalently for both positive and negative stimuli; (Bestelmeyer et al., 2017; Kim et al., 2020; Lewis et al., 2006; Viinikainen et al., 2010, 2012; Winston et al., 2003, 2005). This pattern is typically observed in studies employing conjunction analyses, or when valence is parameterized as intensity (i.e., the absolute value of bipolar valence) or modeled with a quadratic term, yielding a U-shaped or V-shaped relationship between BOLD signal and valence (**Figure 1A**). This pattern has been observed in a distributed network including the OFC, ventromedial prefrontal cortex (vmFPC), dorsomedial prefrontal cortex (dmPFC), anterior cingulate cortex (ACC), middle cingulate cortex (MCC), IFG, STG, MTG, insula, hippocampus, parahippocampal gyrus (PHG), and amygdala (Benelli et al., 2012; Bestelmeyer et al., 2017; Chikazoe et al., 2014; Lewis et al., 2006; Lindquist et al., 2016; Tseng et al., 2016; Viinikainen et al., 2010, 2012). However, an alternative explanation is that increased neural responses to both positive and negative stimuli compared to neutral may not reflect valence but conceptually distinct processes, such as arousal (i.e., the experienced physiological activation, such as being in a calm vs. excited state) or motivational salience (i.e., the significance of stimuli in relation to homeostatic needs and goal-directed attention; Lindquist et al., 2012, 2016; Viinikainen et al., 2012).

Despite extensive research, several methodological issues remain that may hinder a clear understanding of the neural representation of valence. First, most studies have examined valence using a single statistical model, with conclusions typically drawn from patterns of significant activation. However, a significant effect under one model does not exclusively imply that the data provides the best support for the underlying theoretical account. Similarly, it does not preclude alternative models from explaining the same data better when parameterized differently, which may thus support a different theoretical account. As such, inferences about valence may depend critically on the choice of the statistical model rather than the underlying neural organization. Second, applying a single model uniformly across the whole brain implicitly assumes that all regions encode valence in the same manner. It is plausible that different brain regions encode valence using distinct response profiles, for example, some regions may exhibit monotonic responses, whereas others may show valence-general responses (Kim et al., 2020). Ignoring such heterogeneity may obscure region-specific encoding patterns and contribute to inconsistent findings across studies. Only few studies have evaluated multiple alternative modelling approaches within the same dataset (Kim et al., 2017, 2020; Lewis et al., 2006; Lindquist et al., 2016; Viinikainen et al., 2010, 2012), yet even these have primarily relied on the significance of neural activation for each model rather than formal model comparison. This is problematic because a region may show a linear profile when fit with a linear model and a valence-general profile when fit with a quadratic model, making it difficult to determine which one of these alternatives provides the most parsimonious explanation for the observed neural responses.

In addition, much of the existing literature relies on highly controlled stimuli (e.g., emotional images, faces, sounds, and words). While such stimuli provide experimental control and allow researchers to equate potential confounds, they are typically presented as isolated emotional events and therefore do not approximate everyday dynamically-unfolding and highly context-dependent emotional experiences. In contrast, listening to a narrative or watching a movie more closely approximates real-world experience. Neural responses under such naturalistic paradigms have been shown to be more reliable and reproducible than those elicited by controlled stimuli (Hasson et al., 2010; Saarimäki, 2021). Consistent with this, recent studies demonstrate that neural activity within the emotional network of different participants exposed to the same movies or narratives can become synchronized, reflecting shared processing of unfolding emotional dynamics and providing an ecologically valid framework for examining the neural representation of valence across individuals (Nanni-Zepeda et al., 2024; Nummenmaa et al., 2014; Savard et al., 2024; Tan et al., 2022; Vaccaro et al., 2024).

To address these limitations, the present study adopts a formal model comparison framework to characterize how valence-related neural responses are best captured during naturalistic narrative listening. Rather than inferring conclusions from activation patterns under a single model, we directly compare alternative statistical models of valence that differ in their assumed response profiles (e.g., bipolar, valence-general, and bivalent). Model comparison approaches have been widely used in psychology and behavioral sciences but remain underutilized in cognitive neuroscience. Recent developments have enabled Bayesian model comparison within the general linear model (GLM) framework for fMRI data, allowing voxel-wise evaluation of model evidence and selection of the most parsimonious model across participants (Soch et al., 2016, 2017; Soch & Allefeld, 2018). Applying this framework provides a principled approach to dissociate competing statistical accounts of valence representation and to identify potential regional heterogeneity in how valence-related signals are encoded in the brain.

## Method

### Description and preprocessing of fMRI data

We leveraged two datasets from the *Narratives* fMRI data collection (Nastase et al., 2021). Young adults passively listened to four narratives in an MRI scanner. Forty-six participants listened to the narratives *I Knew You Were Black* (*Black*) and *The Man Who Forgot Ray Bradbury* (*Forgot*) in separate sessions, and 19 participants listened to narratives *Slumlord* and *Reach for the Stars One Small Step at a Time* (*Reach*) in single sessions (see Table 1 for the details). One participant was excluded based on the recommendations from the original paper, resulting in a total sample size of 64 people (*M_age_* = 22.67, *SD_age_* = 6.57, 41 females, 23 males).

**Table 1.** Description of fMRI data.

| Narrative | $N$ | $M_{age} (SD)$ | Female/Male |
| --- | --- | --- | --- |
| *Black and Forgot* | 46 | 23.26 (7.55) | 33/13 |
| *Slumlord and Reach* | 18 | 21.17 (2.41) | 8/10 |

We used the preprocessed fMRI data provided in the *Narratives* dataset, which was collected using a 3T scanner with a 1.5s repetition time. For the initial preprocessing steps, the raw fMRI data were temporally and spatially corrected, then normalized to MNI space using fMRIPrep 20.0.5 (Esteban et al., 2019, 2020). The 3dBlurToFWHM program in AFNI 21.1.01 (Cox, 1996) was used to apply a 6mm FWHM Gaussian filter to smooth the data. Confounding variables, including head motion, signals in cerebrospinal fluid and white matter, and time drifts were regressed out (Nastase et al., 2021). The residuals were used as the “clean” fMRI data for our analyses. We resliced all preprocessed images to a matching spatial resolution of 3×3×4 mm^3^ voxels using the 3dResample program in AFNI.

### Subjective Ratings

To capture the variation in valence within narratives and increase the sampling rate of subjective ratings, each sentence was divided into segments following general grammatical rules. Two research assistants independently segmented the stories, extracted the onset and offset timestamps for each segment from the audio recordings, and then conferred to ensure that each segment adhered to the segmentation guidelines, conveyed complete meanings, and followed natural transitions. The raw audio recordings were split into separate audio clips which were then used as stimuli in the behavioral rating experiment.

To obtain the subjective valence and arousal ratings for each segment, four online experiments were conducted with separate samples (total sample size across experiments: *N* = 175; see **Table 2**). Participants were first presented with instructions and examples of how to use a valence-by-arousal grid (**Figure 1A**) to rate their feelings during narrative listening. Then, the audio clips for each segment of the narrative were automatically played, one at a time. A rating screen was shown immediately after each narrative segment, with the instruction “*please rate how you felt about the stimulus*”. All segments were played in their original sequence in the narrative. After completing the listening and rating tasks, participants answered four to five multiple-choice comprehension questions (depending on the narrative) to assess understanding and engagement. Nineteen participants were excluded due to low comprehension accuracy (less than 60%), resulting in a final sample of 156 participants across the four experiments (*M_age_* = 20.75, *SD_age_* = 4.01, 132 females, 24 males).

**Table 2.** Description of subjective rating data.

| Narratives |  |  |  | Participants |  |
| --- | --- | --- | --- | --- | --- |
| Narrative | Duration | Number of segments | $N$ | $M_{age}$ ($SD$) | Female/Male |
| *Black* | 13:20 | 197 | 38 | 20.50 (2.58) | 34/4 |
| *Forgot* | 13:57 | 233 | 39 | 21.00 (4.02) | 28/11 |
| *Slumlord* | 15:03 | 272 | 40 | 20.60 (3.43) | 35/5 |
| *Reach* | 13:45 | 205 | 39 | 20.85 (5.58) | 35/4 |

To evaluate interrater reliability of valence and arousal ratings, we computed intraclass correlation coefficients (ICC) using a two-way random-effects model with absolute agreement, separately for each narrative. We report both single-measure ICC2 and average-measures ICC2k. The former reflects the reliability of an individual rater, whereas the latter reflects the reliability of ratings aggregated across raters. In addition, for each narrative, we computed the Pearson correlation between each participant’s ratings and normative ratings derived from the remaining participants (leave-one-participant-out). For this analysis, normative ratings were computed using a simple (un-weighted) mean across participants. This measure provides a direct estimate of how closely the group-level normative ratings correspond to individual subjective experience.

Although only participants demonstrating sufficient narrative comprehension (accuracy > 60%) were included, substantial individual differences in valence and arousal perception remained (**Figure 1E** and **Table 3**). To obtain representative valence and arousal ratings at the group level, we used a well-established method to compute the weighted average based on the similarity of the valence and arousal ratings across the participants listening to the same narrative (Abdi et al., 2012). Conceptually, this approach downweighs the influence of participants with idiosyncratic ratings. Specifically, we first standardized and normalized each participant’s valence and arousal ratings. Next, we calculated the Hilbert-Schmidt inner product as a similarity measure for every pair of participants. We then applied a principal component analysis (PCA) based on the RV coefficient (i.e., a multivariate generalization of the squared Pearson correlation coefficient) to the resulting inner product matrix and used its first eigenvector to determine participant weights (for details, see Abdi et al., 2012). The weighted average valence and arousal ratings were used for the subsequent fMRI data analyses.

**Table 3.** Interrater reliability of valence and arousal ratings by narrative.

| Rating | Type | Story | ICC | *F* | *df1* | *df2* | *p* | 95% CI |
| --- | --- | --- | --- | --- | --- | --- | --- | --- |
| Valence | ICC2k | *Black* | 0.95 | 21.73 | 196 | 7252 | < .001 | [0.94 0.96] |
|  |  | *Forgot* | 0.93 | 15.08 | 232 | 8816 | < .001 | [0.91 0.94] |
|  |  | *Slumlord* | 0.93 | 18.33 | 271 | 10569 | < .001 | [0.92 0.94] |
|  |  | *Reach* | 0.96 | 31.24 | 204 | 7752 | < .001 | [0.96 0.97] |
|  | ICC2 | *Black* | 0.34 | 21.73 | 196 | 7252 | < .001 | [0.30 0.39] |
|  |  | *Forgot* | 0.25 | 15.08 | 232 | 8816 | < .001 | [0.21 0.29] |
|  |  | *Slumlord* | 0.25 | 18.33 | 271 | 10569 | < .001 | [0.22 0.30] |
|  |  | *Reach* | 0.41 | 31.24 | 204 | 7752 | < .001 | [0.36 0.46] |
| Arousal | ICC2k | *Black* | 0.83 | 8.96 | 196 | 7252 | < .001 | [0.78 0.86] |
|  |  | *Forgot* | 0.66 | 3.90 | 232 | 8816 | < .001 | [0.59 0.72] |
|  |  | *Slumlord* | 0.90 | 13.75 | 271 | 10569 | < .001 | [0.87 0.92] |
|  |  | *Reach* | 0.86 | 8.84 | 204 | 7752 | < .001 | [0.84 0.89] |
|  | ICC2 | *Black* | 0.11 | 8.96 | 196 | 7252 | < .001 | [0.08 0.14] |
|  |  | *Forgot* | 0.05 | 3.90 | 232 | 8816 | < .001 | [0.04 0.06] |
|  |  | *Slumlord* | 0.18 | 13.75 | 271 | 10569 | < .001 | [0.15 0.22] |
|  |  | *Reach* | 0.14 | 8.84 | 204 | 7752 | < .001 | [0.12 0.17] |

### Model comparison within participants

We performed within-participant voxel-wise GLM regressions in SPM 12 (http://fil.ion.ucl.ac.uk/spm) to examine whether BOLD responses varied as 1) a linear function of valence (Bipolar); 2) a quadratic function of valence (Valence-general, U-shaped); 3) a linear function of absolute-transformed valence (Valence-general, V-shape); or 4) a linear function of positivity and negativity separately (Bivalence). The Bipolar and valence-general (V-shaped and U-shaped) models each included a single boxcar function marking the onset and duration of each narrative segment. The Bivalence model included two boxcar functions: one for positive-valence segments and one for negative-valence segments. All models also included the additional modulators described below (**Figure 2A**).

**Figure 2.**
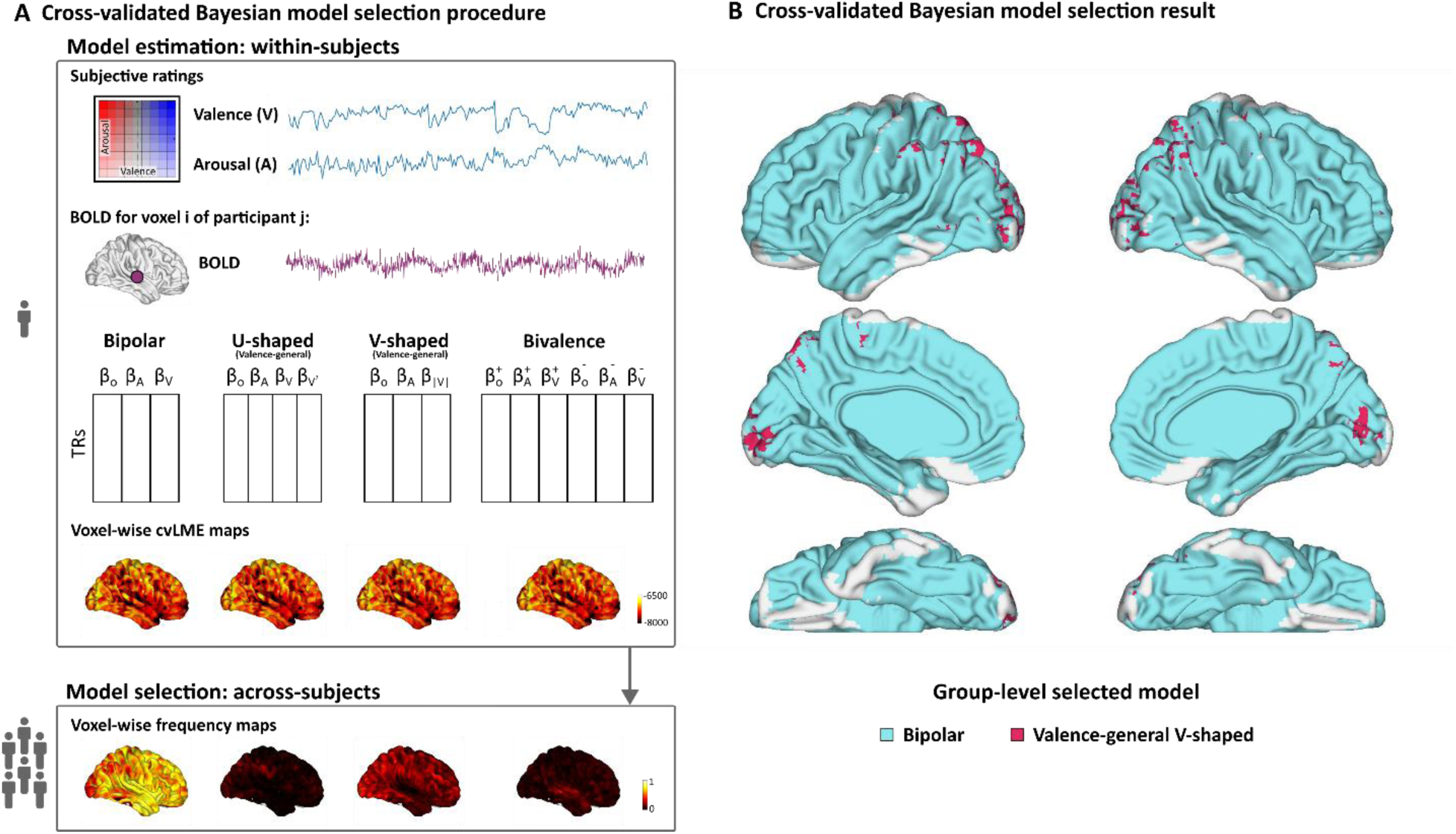
**A.** Schematic representation of the cross-validated Bayesian model selection (cvBMS) process. **B.** Group-level model selection map showing, for each voxel, which model was most often identified as optimal across participants. Voxels supporting the Bipolar model were shown in cool color, whereas those supporting the Valence-general V-shaped model were shown in warm color. The valence-general U-shaped and Bivalence models were not selected for any voxels. Uncolored regions in the orbitofrontal cortex and anterior temporal lobe reflect signal loss across participants. *Notes*. In panel A, for the indices of the modulators, O indicates the boxcar function for the onset, A indicates arousal, and V indicates valence.

#### Bipolar model

This model included two additional parametric modulators: one for arousal and one for valence.

#### Valence-general, U-shaped model

This model included additional linear parametric modulators for arousal and valence as the Bipolar model, plus an additional parametric modulator representing the quadratic (squared) term of the valence ratings to capture U-shaped effects.

#### Valence-general, V-shaped model

This model included an additional linear arousal parametric modulator (as in the Bipolar model), and an absolute-value-transformed valence modulator to model V-shaped effects.

#### Bivalence model

This model was similar to the Bipolar model but modeled positive- and negative-valence segments separately. It included four additional modulators: an arousal parametric modulator and a valence parametric modulator for positive-valence segments, and the same two for negative-valence segments.

Considering that SPM uses a serial orthogonalization procedure which can assign shared variance to earlier parametric modulators and attenuate later modulators, we disabled the default orthogonalization and applied mean-centering to reduce the collinearity among regressors. For all models, arousal ratings were mean-centered prior to modeling. Valence ratings were mean-centered in the Bipolar and Bivalence model, but not in the Valence-general V-shaped and U-shaped models^1^. This was to use neutral valence (rating = 0) as a baseline reference, as required by the definition of the two Valence-general models. All boxcar functions and parametric modulators were convolved with the canonical hemodynamic response function (HRF) to account for the temporal dynamics of BOLD signals. Considering that valence and arousal are known to correlate, we calculated the variance inflation factor (VIF) to examine the multicollinearity among regressors, derived from the *SPM.xX.X* design matrix after convolution. The results showed acceptable multicollinearity for all models (VIF ≤ 5; see **Figure S1-S4**).

To determine which model fits the data best for each voxel, we applied cross-validated Bayesian model comparison (cvBMS) using the SPM-based MACS toolbox (Soch & Allefeld, 2018). For each voxel and each participant, this method calculates a cross-validated version of the log model evidence (cvLME) across fMRI sessions. Specifically, for the narratives *Black* and *Forgot* that were acquired in two separate sessions, one session was used to estimate posterior distributions, which were then used as informative priors for the other session to compute out-of-sample log model evidence. This procedure was repeated across sessions and the resulting values were summed to obtain the cvLME values for each of the four models (Model Family 1). For the narratives *Slumlord* and *Reach* that were acquired in a single session, half-split cross-validation was applied (Soch et al., 2016). These cvLME maps were then converted into posterior model probability maps, and the model with the highest posterior probability was selected as the most optimal model for that voxel and participant (**Figure 2A**).

### Model selection across participants

Because different voxels and participants may favor different models, we applied a between-participant model selection analysis using the MACS toolbox. For each voxel, we applied random-effects Bayesian model selection (RFX BMS), allowing for the possibility that different participants have different optimal models. We computed the likeliest frequency with which each model was selected as the most optimal model over the other three models across participants, resulting in four model frequency maps. We then constructed a between-participants model selection map by assigning each voxel to the model with the highest selection frequency (**Figure 2A**).

### Identifying the voxels modulated by valence

While cvBMS identifies the best-fitting model, it does not indicate whether valence significantly modulates BOLD activity. To address this, we conducted voxel-wise within-participant t-tests (for Bipolar, Valence-general V-shaped, and Bivalence models) and F-tests (for the Valence-general U-shaped model) on the parametric modulators of valence. The F-tests allow us to examine the overall effect of both linear and quadratic terms in the Valence-general U-shaped model. For the Bivalence model, the effects of positive and negative valence were tested separately. Be-tween-participant analyses were then conducted to identify regions where valence modulation effects were consistent across participants. Statistical maps were thresholded using voxel-wise *p* < .001 and cluster-size correction using SPM12. To isolate regions where valence significantly modulated neural activity under the most appropriate model, we binarized the group-level model selection maps and applied these masks to the thresholded group-level statistical maps.

### Localizing the regions broadly activated when listening to narratives

To identify the brain activation responses to narrative listening independently of the affective content, we conducted a group-level analysis based on the effect of the first boxcar function parametric modulator in the Bipolar model. Statistical maps were thresholded as previously described.

### Localizing the regions selectively modulated by arousal after controlling for valence

To examine the unique effect of arousal, we fitted a separate GLM model using the same model as for the Valence Bipolar model, except that we reversed the order in which the arousal and valence regressors were entered into the model, allowing us to examine the linear relationship between BOLD and arousal after removing the variance explained by valence. Group-level analysis was conducted at voxel-wise *p* < .001 with cluster-size correction, similarly to how we examined valence modulation effect for the Bipolar model.

## Conjunction analysis

To isolate regions sensitive only to valence, only to arousal, or to both valence and arousal, we conducted a conjunction analysis. We first generated a binary mask containing the voxels that were significantly correlated with valence after controlling for arousal in the Bipolar model, without differentiating the direction of the correlation. A similar mask was generated for the effect of arousal after controlling for valence. Then, we applied a voxel-wise conjunction analysis to identify the voxels that were broadly sensitive to both valence and arousal, as well as those uniquely responsive to one affective dimension.

## Results

### Normative affective ratings

For valence, interrater reliability of the aggregated ratings was consistently high across narratives (mean ICC2k = 0.94, range = 0.93–0.96). In contrast, reliability at the level of individual raters was low (mean ICC2 = 0.31, range = 0.25–0.41), indicating a robust shared hedonic structure during narrative listening alongside substantial individual variability. For arousal, reliability of the aggregated ratings was moderate to high (mean ICC2k = 0.81, range = 0.66–0.90). In contrast, reliability at the level of individual raters was low (mean ICC2 = 0.12, range = 0.05–0.18), indicating that although a shared arousal structure is present at the group level, individual differences in arousal perception are substantial.

To characterize how well the normative ratings capture individual subjective experience, for each participant, we computed the correlation between their ratings and leave-one-out normative ratings (**Figure 3**). For valence, correlations were moderate to high, with median values ranging from 0.58 to 0.78 across the four narratives. The distributions were relatively tight, indicating strong alignment between individual raters and the shared valence trajectory. For arousal, correlations were lower and more variable than for valence, with median values ranging from 0.26 to 0.52. The distributions were also broader, indicating greater individual differences in arousal perception.

**Figure 3.**
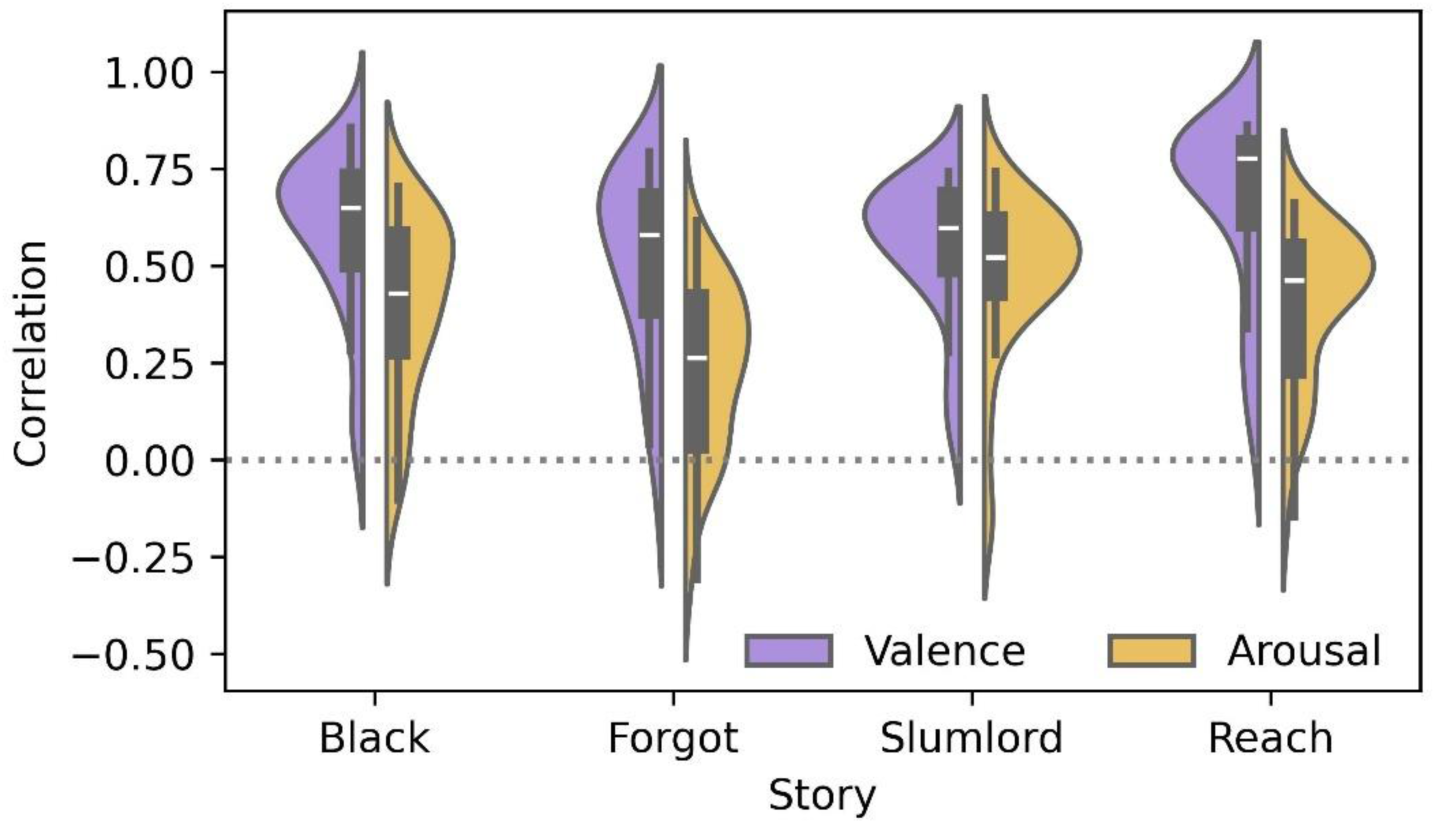
Alignment between individual ratings and leave-one-out normative ratings across narratives. Split violin plots with the superimposed boxplots show the distribution of correlations for valence (blue) and arousal (orange) for each narrative (*Black, Forgot, Slumlord,* and *Reach*). Across narratives, valence generally exhibits higher and more consistent correlations than arousal, with narrative-specific variability.

### Model comparison

The cvBMS results showed that the Bipolar model had the highest model selection frequency across the sample compared to the other three models for most of the brain voxels (**Figure 2B**). The Valence-general V-shaped model was most frequently selected only in several clusters in the parieto-occipital lobe, including bilateral inferior parietal lobule (IPL), angular gyrus (AG), inferior occipital gyrus (IOG), lateral occipital gyrus (LOG), calcarine sulcus, and precuneus (PreCun). Neither the Valence-general U-shaped nor the Bivalence model was selected as the group-level optimal model for any brain voxels.

Considering that the cvBMS framework selects models based on a trade-off between model accuracy and complexity, we cannot rule out the possibility that the Bivalence model was not selected due to its higher complexity (i.e., six parameters, compared to three parameters in the Bipolar and Valence-general V-shaped models, and four parameters in the Valence-general U-shaped model). To address this concern, we reconstructed the Bivalence model with reduced complexity. Specifically, rather than splitting ratings into mutually exclusive positive- and negative-valence segments, we applied an additional STATIS procedure to derive group-level ratings of positive valence and negative valence, separately. For positive valence, negative ratings were set to zero for each participant prior to applying STATIS to compute a weighted average across participants. For negative valence, positive values were set to zero, the sign of the ratings was reversed, and the same STATIS procedure was applied to obtain synthesized negative valence ratings. This approach yielded continuous estimates of both positive and negative valence for each segment. Using these reconstructed regressors, we defined a reduced-complexity version of the Bivalence model with four parameters, including onset, arousal, positive valence, and negative valence regressors^2^. This model was combined with the original three models (Bipolar, Valence-general V-shaped, and Valence-general U-shaped) to form a new model comparison family (Model Family 2). Multicollinearity was acceptable for the reconstructed model (VIF ≤ 5; see **Figure S5**). This additional analysis yielded results largely consistent with those from the original model comparison family (Model Family 1), in which the Bivalence model included six parameters. The group-level model selection map showed that voxels supporting the Bipolar and Valence-general V-shaped models were qualitatively similar to those identified in the original model comparison (**Figure S6A**). No voxels supported the Valence-general U-shaped model. Two clusters in bilateral primary auditory cortex supported the four-parameter Bivalence model.

### Brain activity during narrative listening

The cvBMS analyses identified the best-fitting model for each voxel but did not indicate which specific regressors modulated neural activity. To address this, we next examined neural responses associated with individual regressors. We first examined the effect of narrative listening using the onset time regressor from the Bipolar model, which was selected as the optimal model for most voxels.

In response to narrative listening, increased neural activation was observed in regions associated with speech processing and comprehension, including bilateral STG, STS, MTG, inferior temporal gyrus (ITG), PHG, and left IFG extending to inferior frontal junction (IFJ); reduced neural activation was observed in regions associated with the default mode network (DMN), including bilateral dmPFC, ACC, MCC, STG, SMG/angular gyrus (AG), insula, PreCun, and right IFG (**Figure 4A** and **Table 4**).

**Figure 4.**
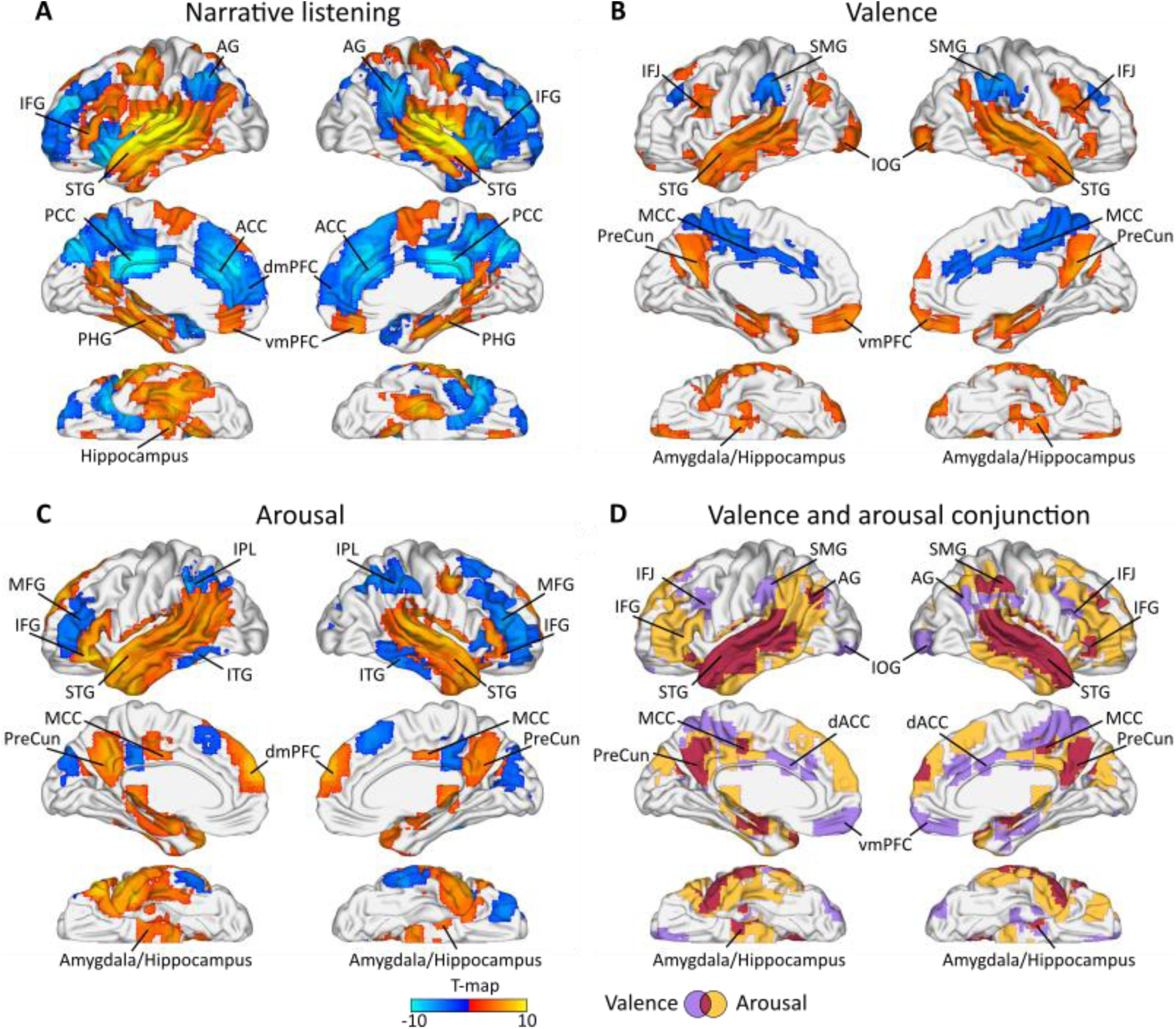
**A-C.** Positive (warm colors) and negative (cool colors) correlation between neural activity and narrative listening (**A**), bipolar valence after controlling for the effect of arousal (**B**), and arousal after controlling for the effect of bipolar valence (**C**). **D.** Conjunction between the unique effect of valence and the unique effect of arousal, without differentiating positive and negative correlations. *Notes*. All results were superimposed on a standard brain surface template and thresholded using cluster-size based correction, voxel-wise *p* < .001.

**Table 4.**
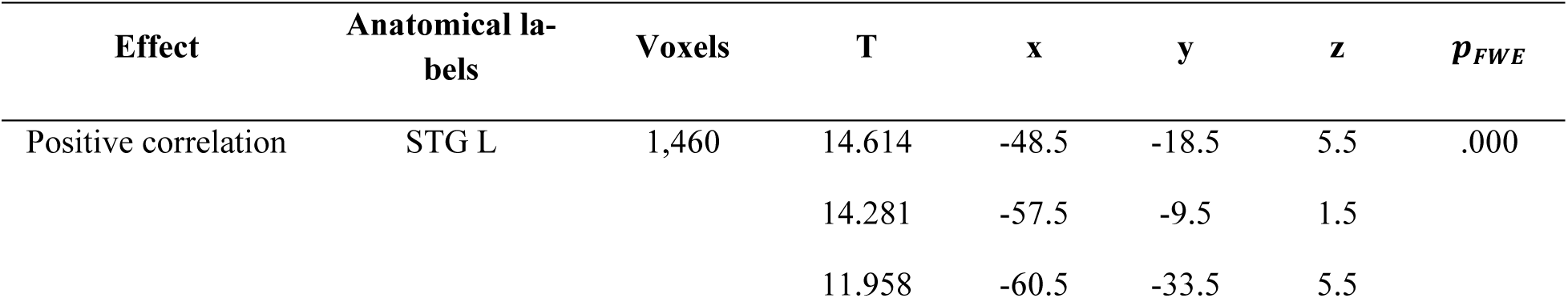
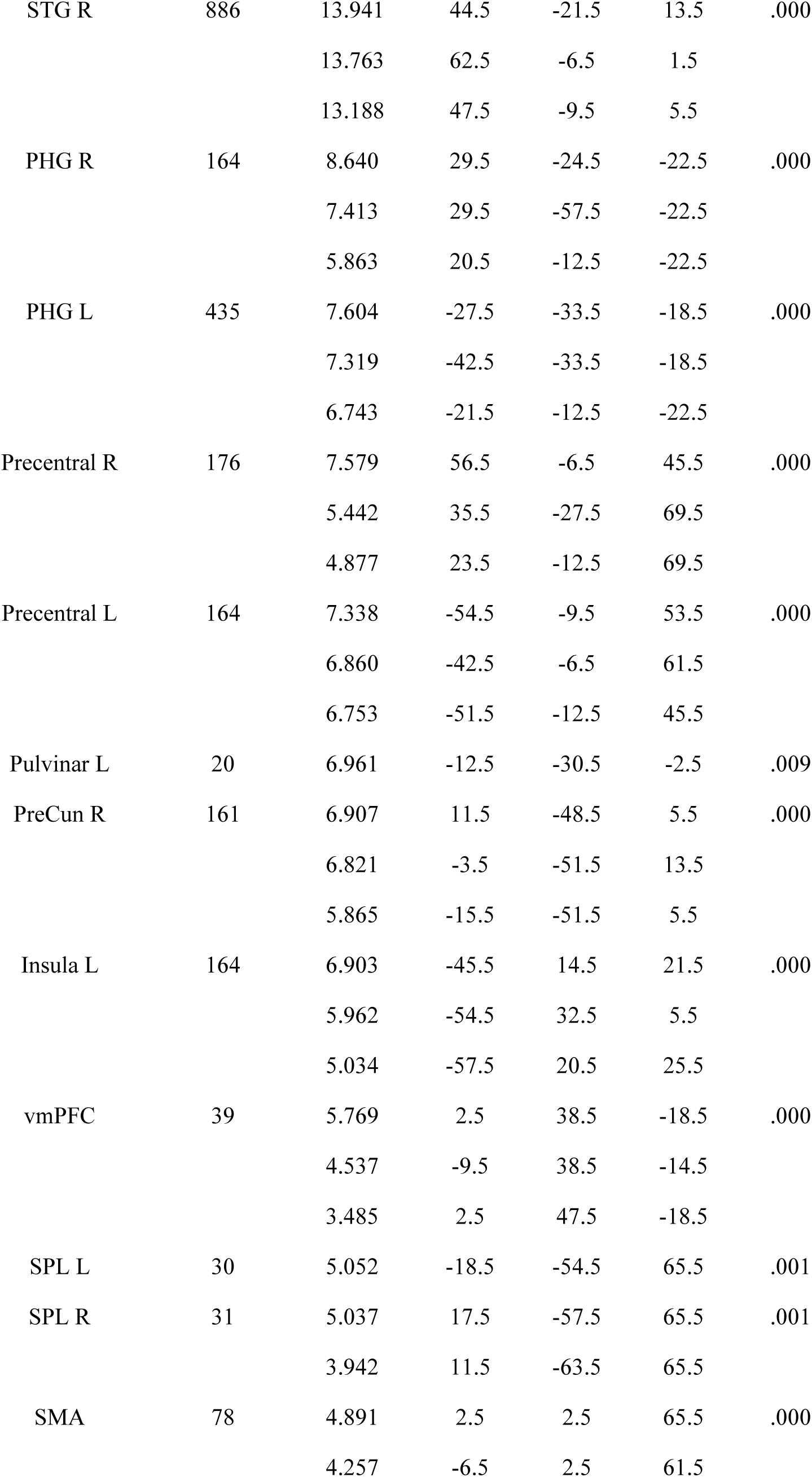
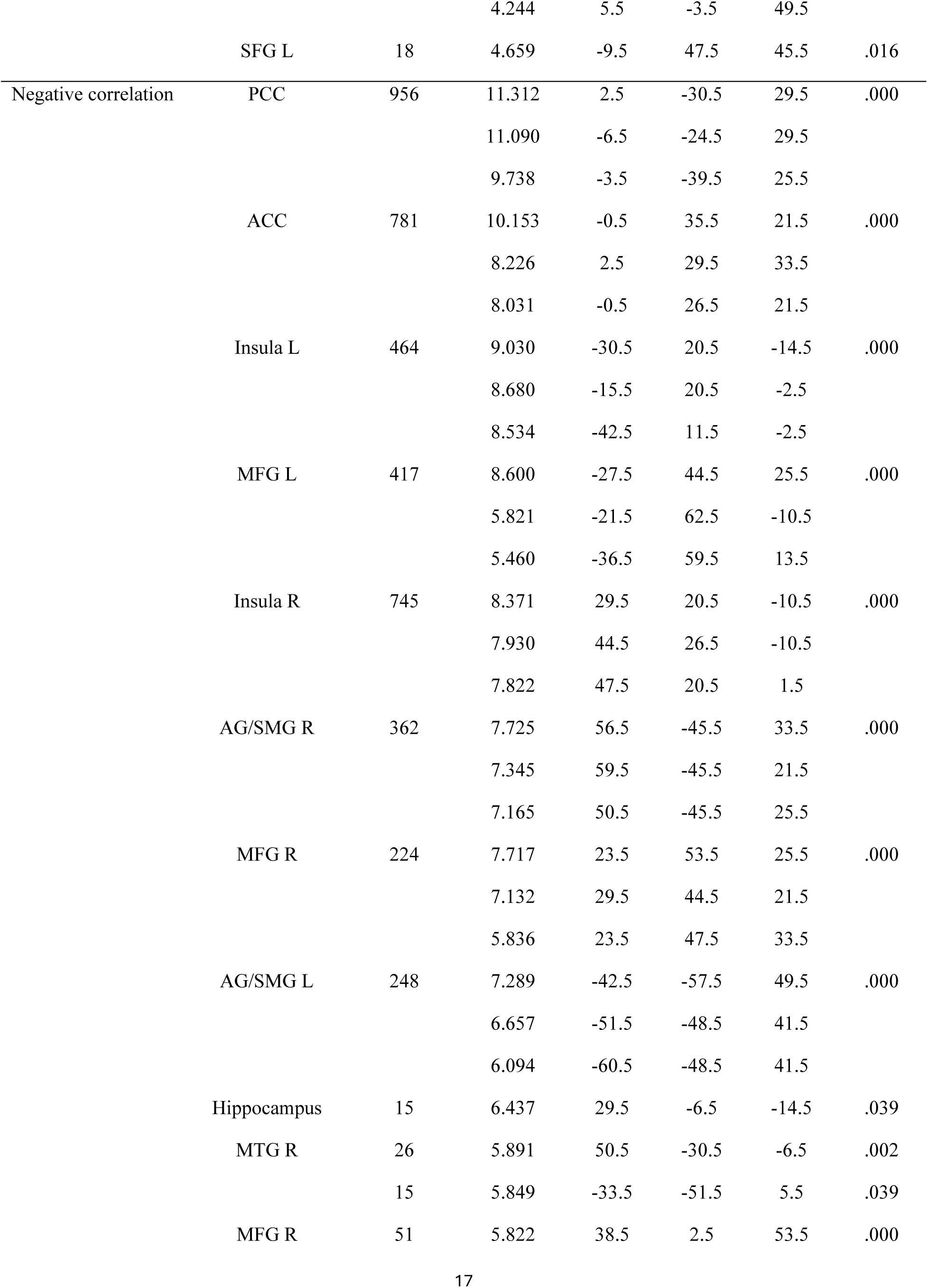
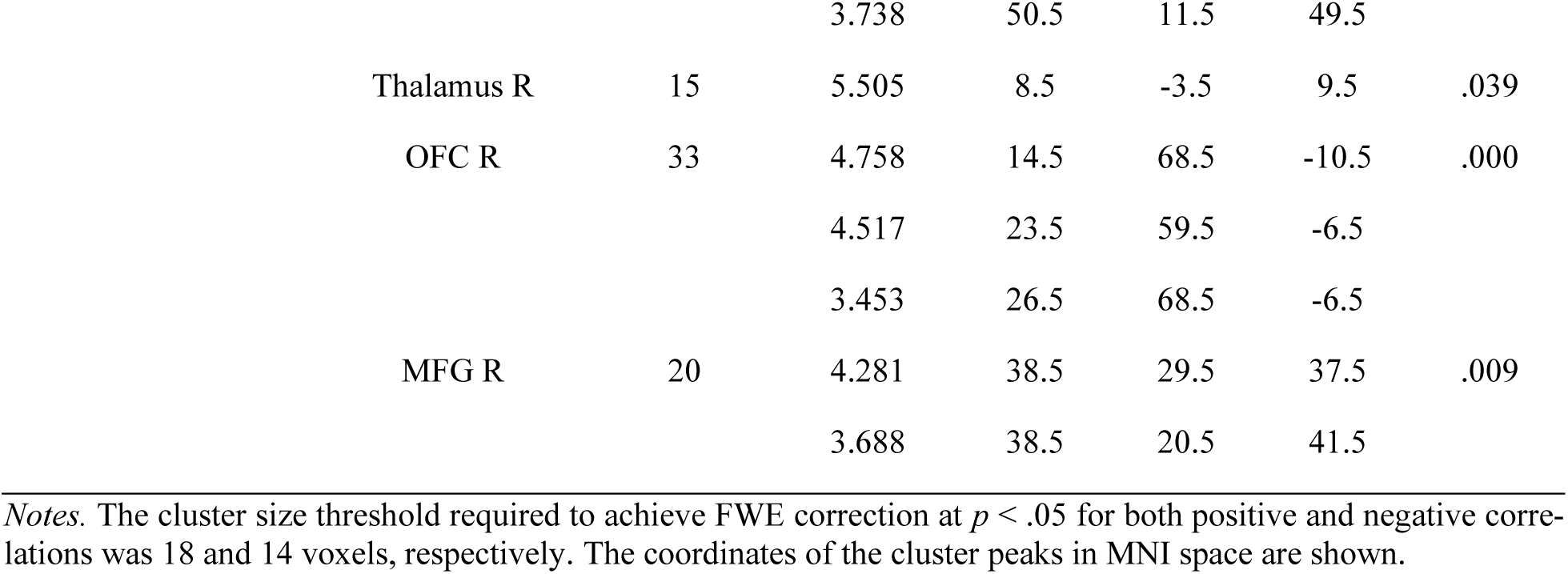
Correlations with narrative listening.

### Modulation effect of valence after controlling for arousal

We conducted a group-level analysis to identify voxels significantly modulated by valence within the regions selected by each model, respectively. In Model Family 1, among the voxels identified by the Bipolar model, after controlling for arousal, valence ratings were significantly positively correlated with brain activation in bilateral vmPFC, IFG/IFJ, STG, STS, AG, PreCun, amygdala, hippocampus, and negatively correlated with brain activation in MFG, SMG, MCC, and PCC (**Figure 4B** and **Table 5**). Among the voxels identified by the Valence-general V-shaped model, none showed significant modulation by valence after controlling for arousal.

**Table 5.**
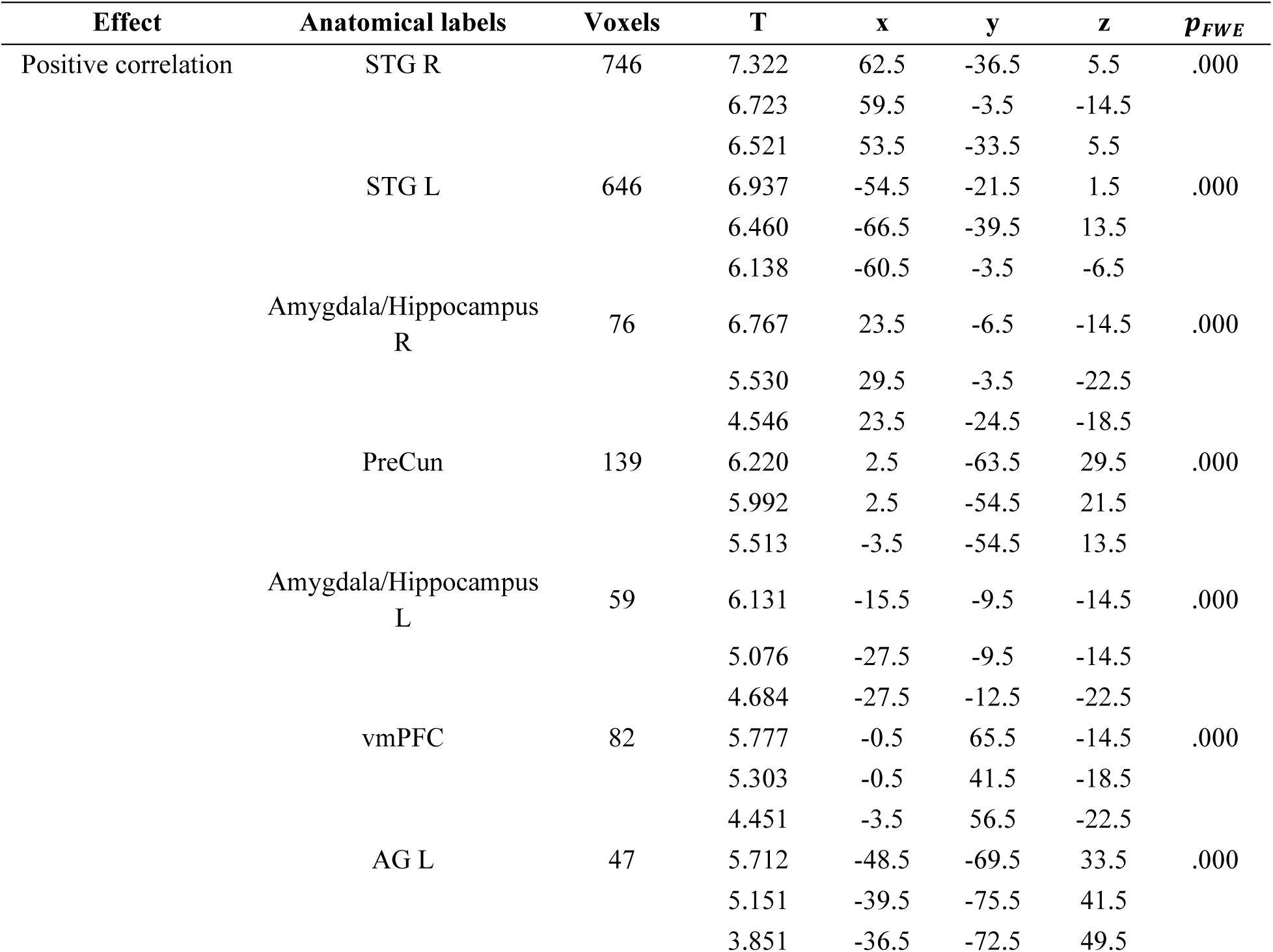
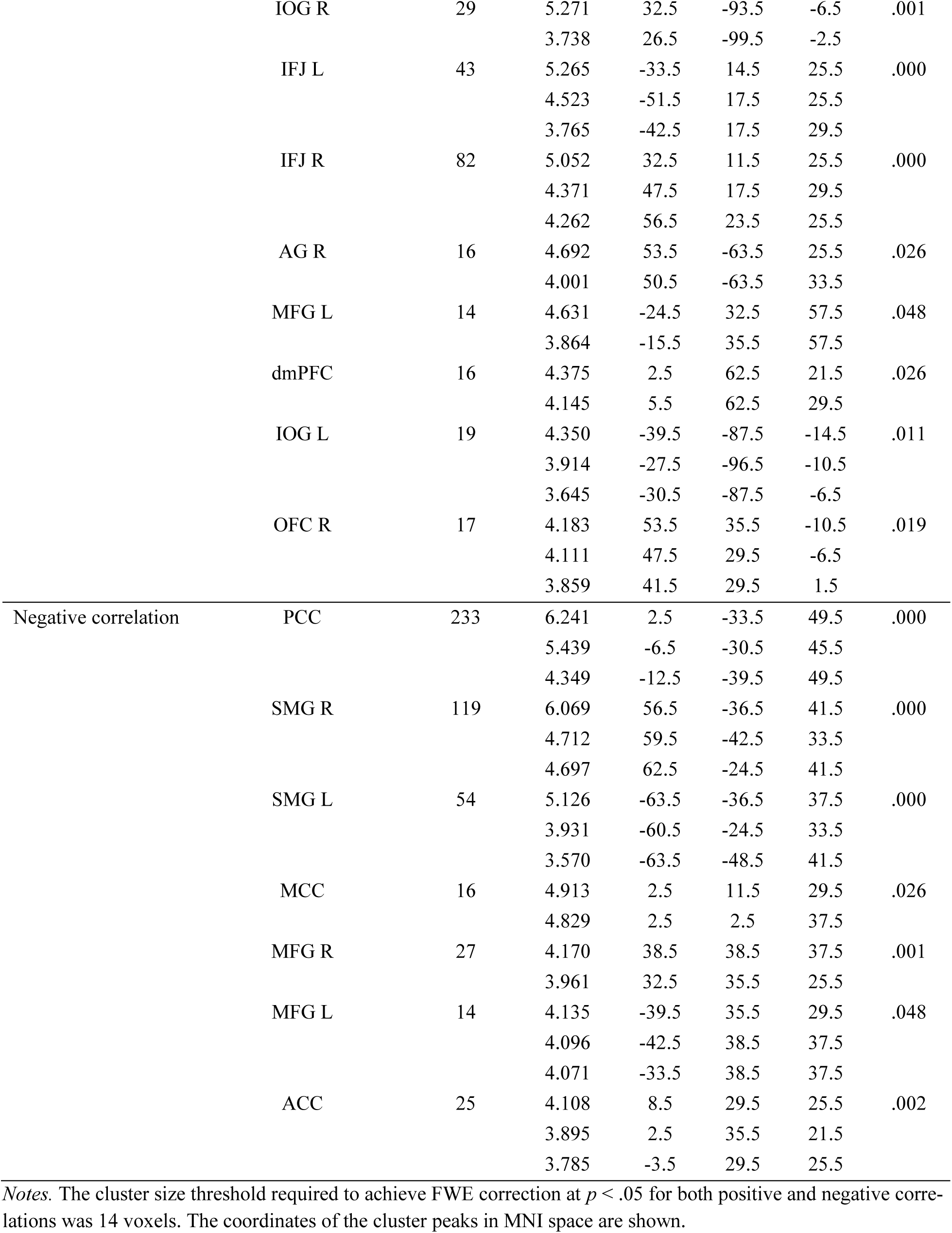
Correlations with bipolar valence after controlling for arousal.

| Effect | Anatomical labels | Voxels | T | x | y | z | $p_{FWE}$ |
| --- | --- | --- | --- | --- | --- | --- | --- |
| Positive correlation | STG R | 746 | 7.322 | 62.5 | -36.5 | 5.5 | .000 |
|  |  |  | 6.723 | 59.5 | -3.5 | -14.5 |  |
|  |  |  | 6.521 | 53.5 | -33.5 | 5.5 |  |
|  | STG L | 646 | 6.937 | -54.5 | -21.5 | 1.5 | .000 |
|  |  |  | 6.460 | -66.5 | -39.5 | 13.5 |  |
|  |  |  | 6.138 | -60.5 | -3.5 | -6.5 |  |
|  | Amygdala/Hippocampus R | 76 | 6.767 | 23.5 | -6.5 | -14.5 | .000 |
|  |  |  | 5.530 | 29.5 | -3.5 | -22.5 |  |
|  |  |  | 4.546 | 23.5 | -24.5 | -18.5 |  |
|  | PreCun | 139 | 6.220 | 2.5 | -63.5 | 29.5 | .000 |
|  |  |  | 5.992 | 2.5 | -54.5 | 21.5 |  |
|  |  |  | 5.513 | -3.5 | -54.5 | 13.5 |  |
|  | Amygdala/Hippocampus L | 59 | 6.131 | -15.5 | -9.5 | -14.5 | .000 |
|  |  |  | 5.076 | -27.5 | -9.5 | -14.5 |  |
|  |  |  | 4.684 | -27.5 | -12.5 | -22.5 |  |
|  | vmPFC | 82 | 5.777 | -0.5 | 65.5 | -14.5 | .000 |
|  |  |  | 5.303 | -0.5 | 41.5 | -18.5 |  |
|  |  |  | 4.451 | -3.5 | 56.5 | -22.5 |  |
|  | AG L | 47 | 5.712 | -48.5 | -69.5 | 33.5 | .000 |
|  |  |  | 5.151 | -39.5 | -75.5 | 41.5 |  |
|  |  |  | 3.851 | -36.5 | -72.5 | 49.5 |  |
|  | I/OG R | 29 | 5.271 | 32.5 | -93.5 | -6.5 | .001 |
|  |  |  | 3.738 | 26.5 | -99.5 | -2.5 |  |
|  | IFJ L | 43 | 5.265 | -33.5 | 14.5 | 25.5 | .000 |
|  |  |  | 4.523 | -51.5 | 17.5 | 25.5 |  |
|  |  |  | 3.765 | -42.5 | 17.5 | 29.5 |  |
|  | IFJ R | 82 | 5.052 | 32.5 | 11.5 | 25.5 | .000 |
|  |  |  | 4.371 | 47.5 | 17.5 | 29.5 |  |
|  |  |  | 4.262 | 56.5 | 23.5 | 25.5 |  |
|  | AG R | 16 | 4.692 | 53.5 | -63.5 | 25.5 | .026 |
|  |  |  | 4.001 | 50.5 | -63.5 | 33.5 |  |
|  | MFG L | 14 | 4.631 | -24.5 | 32.5 | 57.5 | .048 |
|  |  |  | 3.864 | -15.5 | 35.5 | 57.5 |  |
|  | dmPFC | 16 | 4.375 | 2.5 | 62.5 | 21.5 | .026 |
|  |  |  | 4.145 | 5.5 | 62.5 | 29.5 |  |
|  | I/OG L | 19 | 4.350 | -39.5 | -87.5 | -14.5 | .011 |
|  |  |  | 3.914 | -27.5 | -96.5 | -10.5 |  |
|  |  |  | 3.645 | -30.5 | -87.5 | -6.5 |  |
|  | OFC R | 17 | 4.183 | 53.5 | 35.5 | -10.5 | .019 |
|  |  |  | 4.111 | 47.5 | 29.5 | -6.5 |  |
|  |  |  | 3.859 | 41.5 | 29.5 | 1.5 |  |
| Negative correlation | PCC | 233 | 6.241 | 2.5 | -33.5 | 49.5 | .000 |
|  |  |  | 5.439 | -6.5 | -30.5 | 45.5 |  |
|  |  |  | 4.349 | -12.5 | -39.5 | 49.5 |  |
|  | SMG R | 119 | 6.069 | 56.5 | -36.5 | 41.5 | .000 |
|  |  |  | 4.712 | 59.5 | -42.5 | 33.5 |  |
|  |  |  | 4.697 | 62.5 | -24.5 | 41.5 |  |
|  | SMG L | 54 | 5.126 | -63.5 | -36.5 | 37.5 | .000 |
|  |  |  | 3.931 | -60.5 | -24.5 | 33.5 |  |
|  |  |  | 3.570 | -63.5 | -48.5 | 41.5 |  |
|  | MCC | 16 | 4.913 | 2.5 | 11.5 | 29.5 | .026 |
|  |  |  | 4.829 | 2.5 | 2.5 | 37.5 |  |
|  | MFG R | 27 | 4.170 | 38.5 | 38.5 | 37.5 | .001 |
|  |  |  | 3.961 | 32.5 | 35.5 | 25.5 |  |
|  | MFG L | 14 | 4.135 | -39.5 | 35.5 | 29.5 | .048 |
|  |  |  | 4.096 | -42.5 | 38.5 | 37.5 |  |
|  |  |  | 4.071 | -33.5 | 38.5 | 37.5 |  |
|  | ACC | 25 | 4.108 | 8.5 | 29.5 | 25.5 | .002 |
|  |  |  | 3.895 | 2.5 | 35.5 | 21.5 |  |
|  |  |  | 3.785 | -3.5 | 29.5 | 25.5 |  |

Notes. The cluster size threshold required to achieve FWE correction at $p < .05$ for both positive and negative correlations was 14 voxels. The coordinates of the cluster peaks in MNI space are shown.

In Model Family 2, the valence-related modulation observed in the Bipolar model was similar to that in Model Family 1 (**Figure S6B**). Among voxels identified by the V-shaped model, no voxel showed significant modulation by valence after controlling for arousal. Among voxels identified by the four-parameter Bivalence model, both positive and negative valence ratings were negatively correlated with neural activity in bilateral primary auditory cortex after controlling for arousal (**Figure S6C** and **D**), again consistent with the Valence-general account.

### The modulation effect of arousal after controlling for valence

Although we identified most brain regions encoding bipolar valence after statistically controlling for arousal, this does not preclude these regions from also encoding arousal. To examine this possibility, we fitted a new model with the same three regressors as the Bipolar model, but entered the valence regressor first, followed by the arousal regressor. This allowed us to assess the modulation effect of arousal while statistically controlling for valence. We found that arousal was significantly positively correlated with brain activation in bilateral dmPFC, IFG, STG, STS, MTG, PreCun/PCC, MCC, amygdala, and hippocampus, and significantly negatively correlated with activation in bilateral dmPFC, MFG, ITG, MCC, SMG/IPL, and cuneus (Cun) (**Figure 4C** and **Table 6**).

**Table 6.**
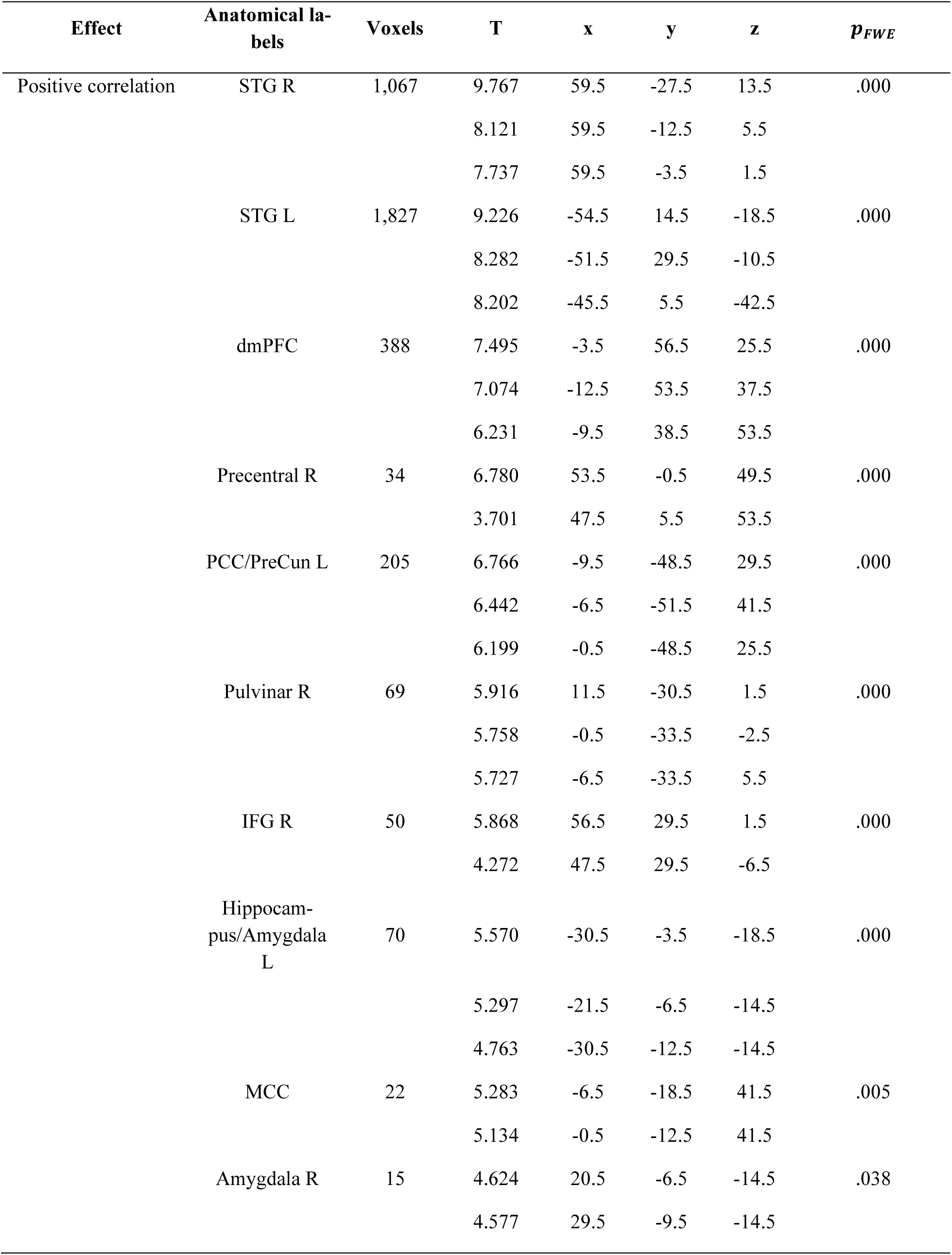
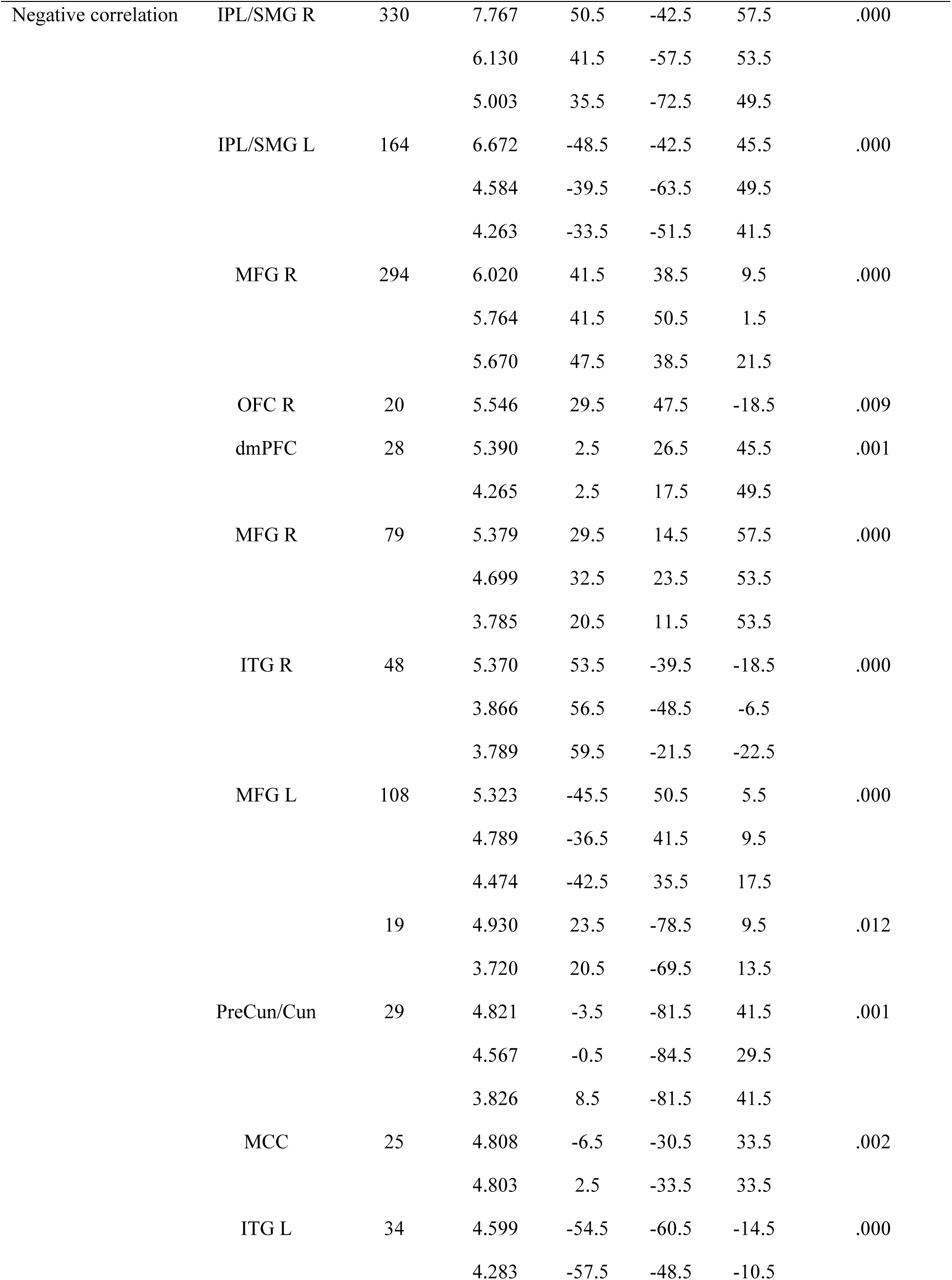
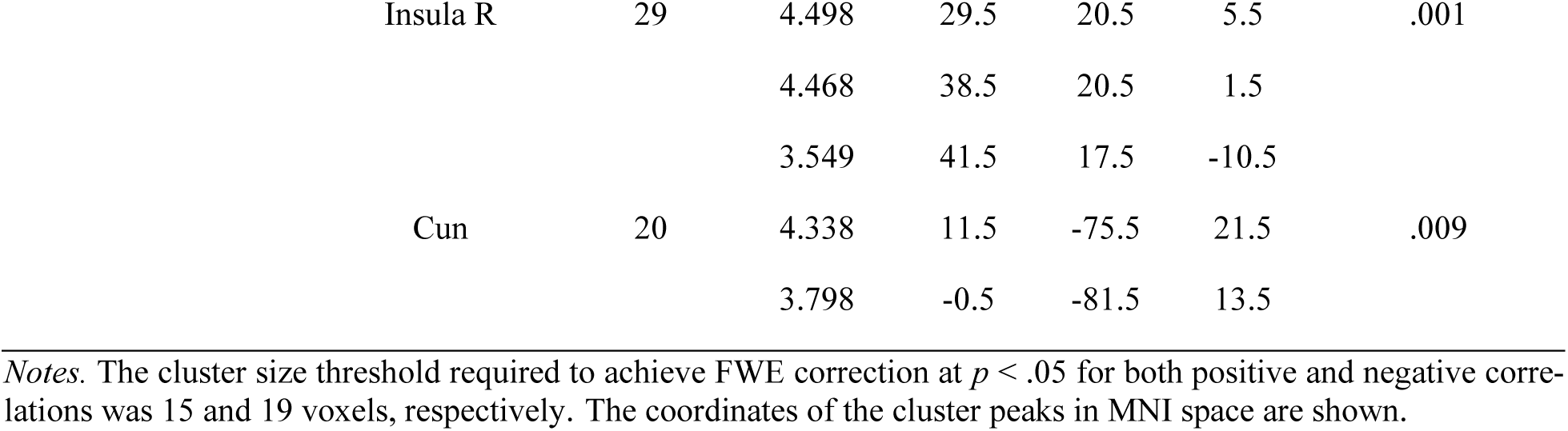
Correlations with arousal after controlling for bipolar valence.

| Effect | Anatomical labels | Voxels | T | x | y | z | $p_{FWE}$ |
| --- | --- | --- | --- | --- | --- | --- | --- |
| Positive correlation | STG R | 1,067 | 9.767 | 59.5 | -27.5 | 13.5 | .000 |
|  |  |  | 8.121 | 59.5 | -12.5 | 5.5 |  |
|  |  |  | 7.737 | 59.5 | -3.5 | 1.5 |  |
|  | STG L | 1,827 | 9.226 | -54.5 | 14.5 | -18.5 | .000 |
|  |  |  | 8.282 | -51.5 | 29.5 | -10.5 |  |
|  |  |  | 8.202 | -45.5 | 5.5 | -42.5 |  |
|  | dmPFC | 388 | 7.495 | -3.5 | 56.5 | 25.5 | .000 |
|  |  |  | 7.074 | -12.5 | 53.5 | 37.5 |  |
|  |  |  | 6.231 | -9.5 | 38.5 | 53.5 |  |
|  | Precentral R | 34 | 6.780 | 53.5 | -0.5 | 49.5 | .000 |
|  |  |  | 3.701 | 47.5 | 5.5 | 53.5 |  |
|  | PCC/PreCun L | 205 | 6.766 | -9.5 | -48.5 | 29.5 | .000 |
|  |  |  | 6.442 | -6.5 | -51.5 | 41.5 |  |
|  |  |  | 6.199 | -0.5 | -48.5 | 25.5 |  |
|  | Pulvinar R | 69 | 5.916 | 11.5 | -30.5 | 1.5 | .000 |
|  |  |  | 5.758 | -0.5 | -33.5 | -2.5 |  |
|  |  |  | 5.727 | -6.5 | -33.5 | 5.5 |  |
|  | IFG R | 50 | 5.868 | 56.5 | 29.5 | 1.5 | .000 |
|  |  |  | 4.272 | 47.5 | 29.5 | -6.5 |  |
|  | Hippocampus/Amygdala L | 70 | 5.570 | -30.5 | -3.5 | -18.5 | .000 |
|  |  |  | 5.297 | -21.5 | -6.5 | -14.5 |  |
|  |  |  | 4.763 | -30.5 | -12.5 | -14.5 |  |
|  | MCC | 22 | 5.283 | -6.5 | -18.5 | 41.5 | .005 |
|  |  |  | 5.134 | -0.5 | -12.5 | 41.5 |  |
|  | Amygdala R | 15 | 4.624 | 20.5 | -6.5 | -14.5 | .038 |
|  |  |  | 4.577 | 29.5 | -9.5 | -14.5 |  |
| Negative correlation | IPL/SMG R | 330 | 7.767 | 50.5 | -42.5 | 57.5 | .000 |
|  |  |  | 6.130 | 41.5 | -57.5 | 53.5 |  |
|  |  |  | 5.003 | 35.5 | -72.5 | 49.5 |  |
|  | IPL/SMG L | 164 | 6.672 | -48.5 | -42.5 | 45.5 | .000 |
|  |  |  | 4.584 | -39.5 | -63.5 | 49.5 |  |
|  |  |  | 4.263 | -33.5 | -51.5 | 41.5 |  |
|  | MFG R | 294 | 6.020 | 41.5 | 38.5 | 9.5 | .000 |
|  |  |  | 5.764 | 41.5 | 50.5 | 1.5 |  |
|  |  |  | 5.670 | 47.5 | 38.5 | 21.5 |  |
|  | OFC R | 20 | 5.546 | 29.5 | 47.5 | -18.5 | .009 |
|  | dmPFC | 28 | 5.390 | 2.5 | 26.5 | 45.5 | .001 |
|  |  |  | 4.265 | 2.5 | 17.5 | 49.5 |  |
|  | MFG R | 79 | 5.379 | 29.5 | 14.5 | 57.5 | .000 |
|  |  |  | 4.699 | 32.5 | 23.5 | 53.5 |  |
|  |  |  | 3.785 | 20.5 | 11.5 | 53.5 |  |
|  | ITG R | 48 | 5.370 | 53.5 | -39.5 | -18.5 | .000 |
|  |  |  | 3.866 | 56.5 | -48.5 | -6.5 |  |
|  |  |  | 3.789 | 59.5 | -21.5 | -22.5 |  |
|  | MFG L | 108 | 5.323 | -45.5 | 50.5 | 5.5 | .000 |
|  |  |  | 4.789 | -36.5 | 41.5 | 9.5 |  |
|  |  |  | 4.474 | -42.5 | 35.5 | 17.5 |  |
|  |  | 19 | 4.930 | 23.5 | -78.5 | 9.5 | .012 |
|  |  |  | 3.720 | 20.5 | -69.5 | 13.5 |  |
|  | PreCun/Cun | 29 | 4.821 | -3.5 | -81.5 | 41.5 | .001 |
|  |  |  | 4.567 | -0.5 | -84.5 | 29.5 |  |
|  |  |  | 3.826 | 8.5 | -81.5 | 41.5 |  |
|  | MCC | 25 | 4.808 | -6.5 | -30.5 | 33.5 | .002 |
|  |  |  | 4.803 | 2.5 | -33.5 | 33.5 |  |
|  | ITG L | 34 | 4.599 | -54.5 | -60.5 | -14.5 | .000 |
|  |  |  | 4.283 | -57.5 | -48.5 | -10.5 |  |

|  |  |  |  |  |  |  |
| --- | --- | --- | --- | --- | --- | --- |
| Insula R | 29 | 4.498 | 29.5 | 20.5 | 5.5 | .001 |
|  |  | 4.468 | 38.5 | 20.5 | 1.5 |  |
|  |  | 3.549 | 41.5 | 17.5 | -10.5 |  |
| Cun | 20 | 4.338 | 11.5 | -75.5 | 21.5 | .009 |
|  |  | 3.798 | -0.5 | -81.5 | 13.5 |  |

*Notes.* The cluster size threshold required to achieve FWE correction at $p < .05$ for both positive and negative correlations was 15 and 19 voxels, respectively. The coordinates of the cluster peaks in MNI space are shown.

### Conjunction between the effect of valence and arousal

To isolate the brain regions that are broadly sensitive to both valence and arousal and that are uniquely sensitive to one dimension, we computed a conjunction map (**Figure 4D**) between the voxels modulated by valence after the variance of arousal was removed, and the voxels modulated by arousal after the variance of valence was removed, without differentiating positive or negative correlation. We found that bilateral STG, STS, MCC, PreCun, amygdala/hippocampus, left AG, right SMG, right MFG, and right dmPFC were sensitive to both valence and arousal. In contrast, regions including bilateral vmPFC, IFG/IFJ, dorsal ACC (dACC), MCC, and IOG, and left SMG were specifically sensitive to valence. Regions including bilateral dmPFC, IFG, and PHG were specifically sensitive to arousal.

## Discussion

Our study combined a formal Bayesian model comparison framework with parametric modulation models to examine how hedonic valence is represented in the brain under an ecologically valid context during narrative listening. By comparing multiple statistical models of valence within the same dataset, we evaluated which model provides the best fit for characterizing affect-related neural responses. Across most brain voxels and participants, a linear model corresponding to the bipolar representation of valence was consistently selected as the optimal model.

It is important to distinguish between model selection and statistical inference on individual regressors. The cross-validated Bayesian model comparison framework identifies the model that best explains the overall BOLD signal in each voxel, taking into account both goodness-of-fit and model complexity, weighing on the tradeoff between prediction accuracy and overfitting. However, selection of a given model does not necessarily imply that the parametric modulator of interest (e.g., valence) exhibits a statistically significant effect. Alternative models, including reparameterizing bipolar valence as “valence intensity” (Valence-general V-shaped), using additional quadratic term to delineate the non-linear shape (Valence-general U-shaped), and reconstruct bipolar valence as bivalent scales (Bivalence), were selected in some voxels when compared within different model families. However, when we further examined the neural activity modulated by valence, those voxels showed no or little correspondence with valence. This suggests that although these models captured more variance, the voxels did not reliably encode valence in the hypothesized manner. Importantly, for the voxels selected by the four-parameter Bivalence model in bilateral Heschl’s gyrus, their BOLD signals significantly correlated with both positive and negative valence, largely resembling a valence-general pattern in primary auditory cortex associated with auditory processing in the narrative listening task, rather than reflecting independent encoding of positive and negative valence per se.

Interestingly, our findings supporting the Bipolar representation of valence contrast with previous studies directly comparing statistical models within the same data, which have more often supported the Valence-general or Bivalence hypotheses. However, key methodological differences may explain these discrepancies. First, prior studies often relied on activation-based approaches, interpreting any significant activation as support for a given model. However, statistical significance alone does not indicate good model fit, nor does its absence imply poor fit. Null results for the Bipolar model in earlier work may stem from low statistical power (N < 20) rather than the absence of a true effect. In our data, the neural activity in bilateral STG/STS was significantly modulated by both bipolar valence (Bi-polar model) and absolute-transformed valence (Valence-general V-shaped model), creating ambiguity about the underlying valence–BOLD relationship. However, model comparison using cvBMS showed that the Bipolar model provided a significantly better fit than the V-shaped model for most participants at the group level. This demonstrates that cvBMS can resolve such ambiguity and underscores the importance of jointly considering overall model fit and the statistical significance of the regressor of interest.

Second, previous studies often used highly controlled emotional words (Lewis et al., 2006), static pictures (Viinikainen et al., 2010) and sounds (Viinikainen et al., 2012) or meta-analysis based on studies using factorial designs that treat valence as categorical variables (Lindquist et al., 2016). The rapid presentation of isolated static stimuli may not reflect how emotions unfold in real-life. The observed brain activation in response to both positive and negative stimuli in these studies may alternatively reflect difference between emotional and neutral stimuli driven by salience in a laboratory setting, rather than affective processing representative of day-to-day life experiences. Also, treating valence as categorical variables (e.g., contrasting positive vs. neutral or negative vs. neutral conditions) may not capture the full range of BOLD-valence relationships, considering that valence is conceptualized as a continuous spectrum. In contrast, in our study, examining how BOLD is modulated by continuous valence ratings during immersive narrative listening allows affective states to unfold more dynamically, providing a more ecologically valid context to examine the neural representation of valence. Aligning with this assumption and consistent with our findings, using naturalistic movie-watching, Kim et al. (2020) reported engagement of STS, MFG, and mPFC in predicting signed bipolar valence. Similarly, using a narrative-listening paradigm, Wallentin et al. (2011) found valence-related activation in the bilateral MTG, left IFG, left temporoparietal junction, PreCun, and left vmPFC. Although they found no effect for the Bipolar model, this may be attributed to the high familiarity of the narrative (“The Ugly Duckling”) or the method used to extract valence ratings at the sentence level, which may not capture more time-resolved valence changes that typically happen between clauses and within sentences.

Our findings provide novel insights into the neural representation of affective experience during narrative listening. BOLD signal varied linearly with bipolar valence in distributed frontal-temporo-parietal regions, midline regions, as well as subcortical structures like the amygdala and hippocampus. Many of these regions, including the bilateral STG, STS, MCC, amygdala, hippocampus, and PreCun, were also modulated by arousal. While the Circumplex Model of Affect (Russell, 1980) posits valence and arousal as independent affective dimensions, accumulating empirical evidence suggests they are not strictly independent, as highly positive or negative affective states tend to exhibit higher arousal. Prior studies have shown that regions such as the bilateral STG, MTG, and amygdala encode both dimensions (Baucom et al., 2012; Bestelmeyer et al., 2017; Mourão-Miranda, 2003; Wallentin et al., 2011), indicating a convergence between our findings and prior studies using more tightly controlled, short-duration stimuli.

At the same time, the involvement of these regions should not be taken as evidence of valence-specific coding *per se*. In particular, the amygdala has been variably characterized as preferentially responsive to negative valence, broadly responsive across valence, or more fundamentally tuned to affective salience and motivational relevance (Costafreda et al., 2008; Lindquist et al., 2012, 2016; Pessoa, 2010; Sergerie et al., 2008). Consistent with these perspectives, the co-modulation of these regions by both valence and arousal in the present study suggests that their activity may reflect more general processes such as salience and novelty detection. From this perspective, rather than reflecting stable and directional encoding of positive and negative valence, the observed direction of the bipolar valence association in these regions may be context-dependent, shaped by the distribution of salient events within the narrative and the extent to which positive versus negative moments capture attention or motivational relevance.

Focusing on regions uniquely sensitive to valence while controlling for arousal, our findings showing negative correlations with valence ratings largely replicate those reported by Nummenmaa et al. (2014), who used both GLM analyses and a model-free inter-subject correlation approach during a similar narrative listening task. As valence shifted from positive to negative, neural activity increased in bilateral SMG and PCC, and became more temporally synchronized across participants in dACC, MCC, PCC, and somatosensory regions during narrative listening. Similar increases in synchronization associated with negative valence in midline regions have also been observed during movie watching (Nummenmaa et al., 2012). Functionally, the SMG has been implicated in empathy and mentalizing (Mar, 2011; Silani et al., 2013), while the dACC and MCC are key nodes of the salience network, supporting the detection of motivationally relevant stimuli and the coordination of adaptive responses (Seeley et al., 2007). In contrast, the PCC is a core node in the default mode network, associated with episodic, contextual, and internally oriented processing (Fernandino & Binder, 2024; Greicius et al., 2003; Mar, 2011). The engagement of these regions during the processing of increasingly negative information may reflect a distinct cognitive–affective state, in which listeners integrate salient external cues with internally oriented processes to comprehend negative events and infer characters’ distress.

In contrast, regions showing positive correlations with valence ratings provide both replication and extension of prior findings. Consistent with previous work, we observed increased activation in the bilateral vmPFC associated with increased positive valence ratings. This aligns with a meta-analysis demonstrating that the vmPFC uniquely supports the Bipolarity hypothesis, with BOLD signals particularly sensitive to positive valence (Lindquist et al., 2016). Extensive research in humans and animals has identified the medial prefrontal and orbitofrontal cortices as critical hubs for hedonic experience and pleasure coding (Berridge & Kringelbach, 2015; Kringelbach, 2005), highlighting the vmPFC’s role in integrating positive affect and evaluating hedonic stimuli in naturalistic contexts. At the same time, we also observed increased activation associated with higher positive valence ratings in regions not traditionally linked to valence processing, including bilateral IFJ and IOG, typically involved in semantic representation and cognitive control (Ralph-Lambon et al., 2017). Growing evidence suggests these regions may modulate affective experience, particularly in naturalistic settings. For example, Nummenmaa et al. (2014) reported enhanced neural synchronization in IFG and mPFC with higher positive valence, while Thye et al. (2023) found heightened activation in right superior and IOG driven by higher valence at word-level during narrative listening. Similarly, Abdel-Ghaffar et al. (2024) showed that the occipital-temporal cortex encodes both affective content and semantic categories in response to natural images. Emotion regulation studies further show IFG is engaged when actively regulating emotional intensity through cognitive reappraisal (McRae et al., 2012; Otto et al., 2014). Together, these findings suggest that positive affect in naturalistic contexts may involve greater integration with semantic and conceptual processes, potentially supporting the interpretation of affective meaning.

Interestingly, some regions sensitive to valence in our narrative listening task, such as the IFG, PCC, and precuneus, have not been consistently reported in prior studies using highly controlled stimuli. One possible explanation relates to differences in temporal receptive windows (TRWs) across cortical regions. Work using naturalistic paradigms, including narrative listening and movie watching, has shown that higher-order regions such as the default mode network and lateral prefrontal cortex integrate information over longer timescales, accumulating contextual and semantic information across extended periods (Baldassano et al., 2018; Hasson et al., 2008; Lerner et al., 2014). In contrast, paradigms employing brief, isolated stimuli may preferentially engage regions with shorter TRWs that respond to transient affective features. From this perspective, the recruitment of IFG, PCC, and precuneus in the present study may reflect the gradual construction of affective meaning as information unfolds over time, a process that is more readily captured in naturalistic settings.

There are several methodological limitations that warrant further discussion. These limitations not only constrain the interpretation of our findings but also reflect broader, long-standing challenges in affective neuroscience. First, while our model selection approach supports the Bipolarity hypothesis, the use of BOLD signals restricts our analysis to the voxel level. Some evidence suggests that at the neuronal level, valence representation may align more closely with the Bivalence hypothesis (Belova et al., 2008; Morrison & Salzman, 2009; also see Lindquist et al., 2016 for a discussion). Accordingly, our findings do not necessarily contradict the affective workspace hypothesis (Barrett & Bliss-Moreau, 2009; Lindquist et al., 2012, 2016), which proposes that neural populations flexibly participate in representing affective states, such that the same regions may contribute to both positive and negative experiences depending on context. In addition, although we explored multiple ways of specifying the Bivalence model, the frequentist framework (e.g., parametric modulation within a GLM) cannot provide direct support for the Bivalence theoretical account. This limitation arises because the Bivalence account is defined by two criteria: for a voxel to uniquely encode positive valence, (1) BOLD activity should be significantly associated with positivity, and (2) BOLD activity should not be associated with negativity. While standard GLM approaches can test the first criterion by rejecting a null hypothesis, they are not well suited to provide evidence in favor of the second criterion, which requires support for the absence of an effect. Accordingly, future work may benefit from Bayesian frameworks that can more directly evaluate evidence for both the presence and absence of effects.

Second, we used group-level synthesized affective ratings from an independent sample to localize neural correlates of valence during narrative listening. This approach has been successfully applied in prior work using similar naturalistic paradigms (Kim et al., 2020; Wallentin et al., 2011). Although inter-rater reliability was high at the group level (ICC2k > 0.9 for valence and > 0.8 for arousal), we also observed substantial individual variability (ICC2 ≈ 0.3), indicating meaningful differences in subjective affective experience. These individual differences are unlikely to reflect low engagement, as raters with poor comprehension were excluded. Instead, they may be related to prior experience-sampling findings suggesting that individuals can exhibit idiosyncratic profiles of core affect (Barrett & Bliss-Moreau, 2009). Our approach, which focuses on a representative affective experience derived from a weighted average across an independent sample, is inherently limited insofar as it can only capture the variance associated with this shared experience. From a constructionist perspective, such additional variance may reflect affective states shaped by individual-specific factors, including personal experiences, beliefs, and temporally dynamic bodily states. Moreover, our group-level cvBMS framework based on model selection frequency across participants does not allow us to identify whether distinct neural representational profiles track these individual differences.

Third, although narrative listening offers high ecological validity, the correlational nature of our analysis limits causal inference, especially considering that the normative affective ratings may correlate with other confounding features embedded in the narrative stimuli. A related limitation is that our method cannot fully dissociate affective processing from semantic knowledge of valence, sometimes referred to as “affective valence” versus “semantic valence” (Itkes et al., 2017; Itkes & Kron, 2019). This distinction is particularly relevant in a narrative listening context, which necessarily engages auditory and language networks. For example, regions such as the inferior frontal gyrus and angular gyrus, implicated in our results, are also well known for their roles in semantic processing and control (Binder et al., 2009; Ralph-Lambon et al., 2017). More broadly, acoustic, prosodic, semantic, and affective features of naturalistic stimuli are inherently intercorrelated, making it difficult to isolate their unique contributions using the present approach. Addressing this challenge will require methodological frameworks that can better accommodate such correlated, high-dimensional predictors. At the same time, more controlled experimental paradigms also remain valuable for isolating specific mechanisms through targeted manipulation.

Finally, our samples were predominantly young adult females, and prior research suggests affective processing varies by sex and age (MacCormack et al., 2020). Broader sampling is needed to generalize findings and better understand how valence is represented across diverse populations.

## Conclusions

In sum, we present the first study to apply a cross-validated Bayesian model comparison framework within a naturalistic narrative listening paradigm to formally evaluate which statistical model best characterizes valence representation under ecologically valid conditions. Our findings suggest that hedonic valence is best characterized by a bipolar representation for BOLD signals. While valence-related signals were distributed across cortical and subcortical regions, many areas (e.g., amygdala, STG, hippocampus) were jointly modulated by arousal, suggesting sensitivity to affective salience and motivational relevance rather than valence-specific coding, with effects that may vary depending on contextual dynamics within the narrative. In contrast, regions such as vmPFC was associated with positive valence, while regions involved in salience detection and empathy, such as the dACC, MCC, and SMG, were associated with negative valence. Additionally, areas not typically implicated in controlled affective paradigms, including the PCC, precuneus, IFG and IOG, also encoded valence, emphasizing the importance of using ecologically valid, naturalistic stimuli. These results underscore the value of naturalistic paradigms in affective neuroscience and advance our understanding of how affect is represented in the brain during real-world experiences.

## Supporting information

Suupplemental Materials

## Acknowledgments

We thank Dr. Kristen Lindquist for insightful discussions and valuable feedback. We thank Savanah Belt and Andrew Boldy for their help with narrative segmentation.

## Footnotes

1 Mean-centering shifts the intercept (i.e., the onset regressor) and its interpretation but does not alter the estimated slope coefficients (i.e., beta values associated with the modulators). We ran additional analyses comparing the same model with and without mean-centering. As expected, mean-centering substantially reduced multicollinearity between the onset regressor and the modulator regressors.

2 We also conducted an additional analysis in which the Bivalence model was reconstructed as a three-parameter model (onset, positive valence, and negative valence) without controlling for arousal, yielding the same number of parameters as the Bipolar and Valence-general V-shaped models. The group-level cvBMS results were qualitatively similar to those obtained with the four-parameter model that included arousal. Since the exclusion of arousal in this model creates a confounder with other models controlling for the effect of arousal, these results were not included.

