## Supplementary material for "Neural representation of hedonic valence during narrative listening": Suupplemental Materials

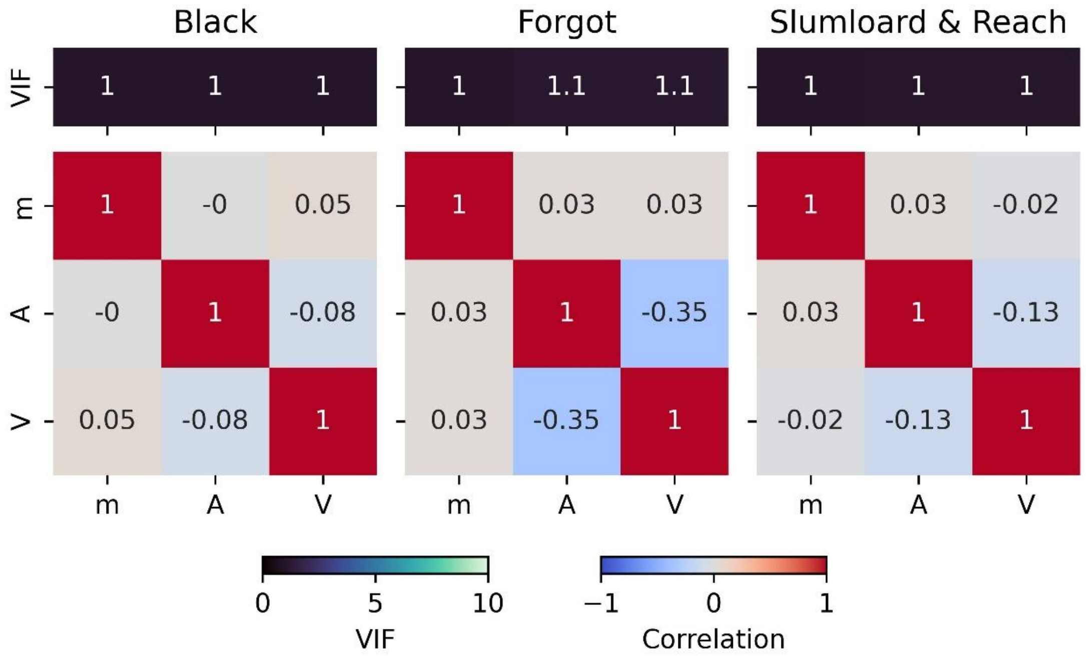


**Figure S1. Variance inflation factors (top) and inter-regressor correlations (bottom) for the bipolar model across stories.** *Notes.* m = the onset time regressor; V = Valence regressor; A = Arousal regressor. Both the valence and arousal ratings were mean-centered first. All parameters were calculated based on the regressors after convolution.

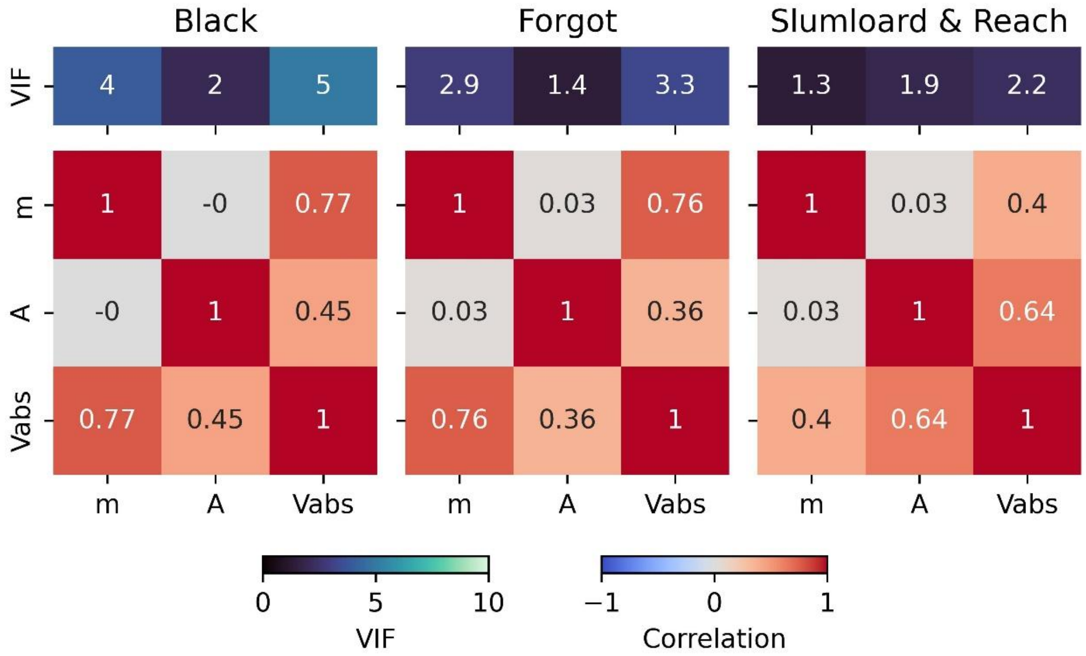


**Figure S2. Variance inflation factors (top) and inter-regressor correlations (bottom) for the valence-general V-shaped model across stories.** *Notes.* m = the onset time regressor; A = the arousal regressor; Vabs = the absolute transformed valence regressor. Only arousal ratings were mean-centered. All regressors were convolved with canonical HRF.

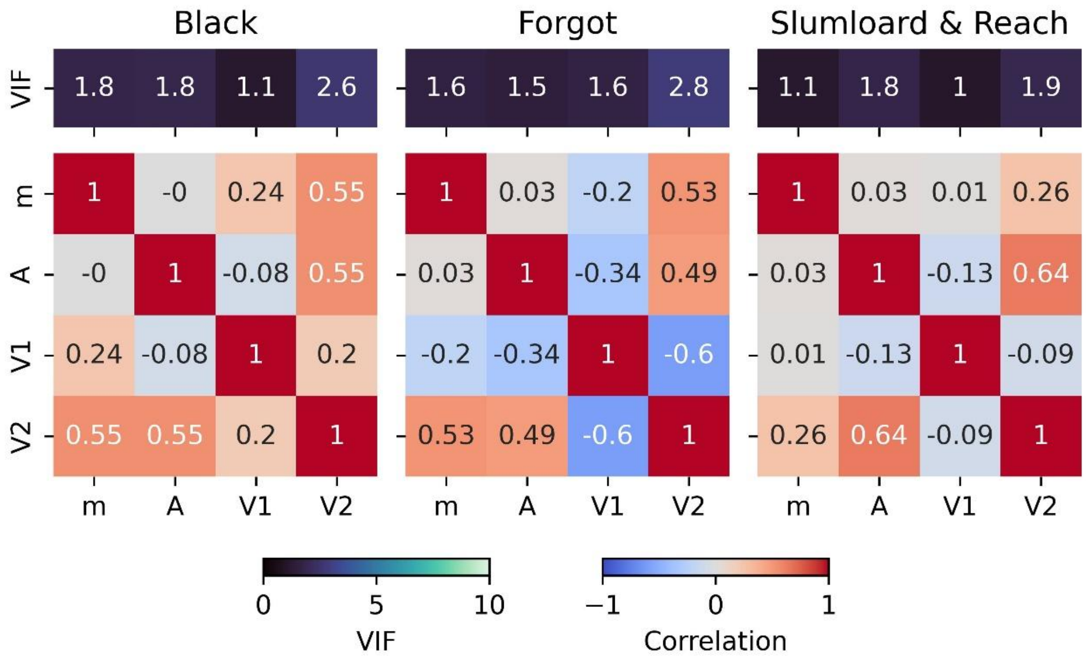


**Figure S3. Variance inflation factors (top) and inter-regressor correlations (bottom) for the valence-general U-shaped model across stories.** *Notes.* m = the onset time regressor; A = the arousal regressor; V1 = the linear valence regressor; V2 = the quadratic valence regressor. Only arousal ratings were mean-centered. All regressors were convolved with canonical HRF.

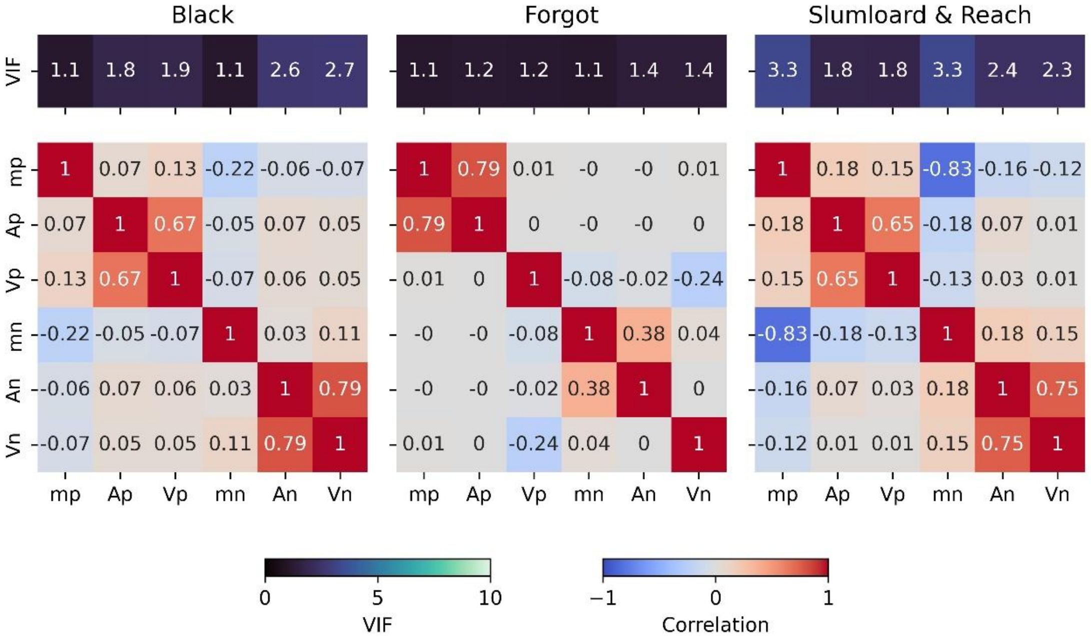


**Figure S4. Variance inflation factors (top) and inter-regressor correlations (bottom) for the six-parameter Bivalence model across stories.** *Notes.* mp = the onset time regressor for positive valence; Ap = the arousal regressor for positive valence; Vp = the valence regressor for positive valence; mn = the onset time regressor for negative valence; An = the arousal regressor for negative valence; Vn = the valence regressor for negative valence. Both valence and arousal ratings were mean-centered first. Valence and arousal ratings were computed using a single Rv-PCA procedure. All regressors were convolved with canonical HRF.

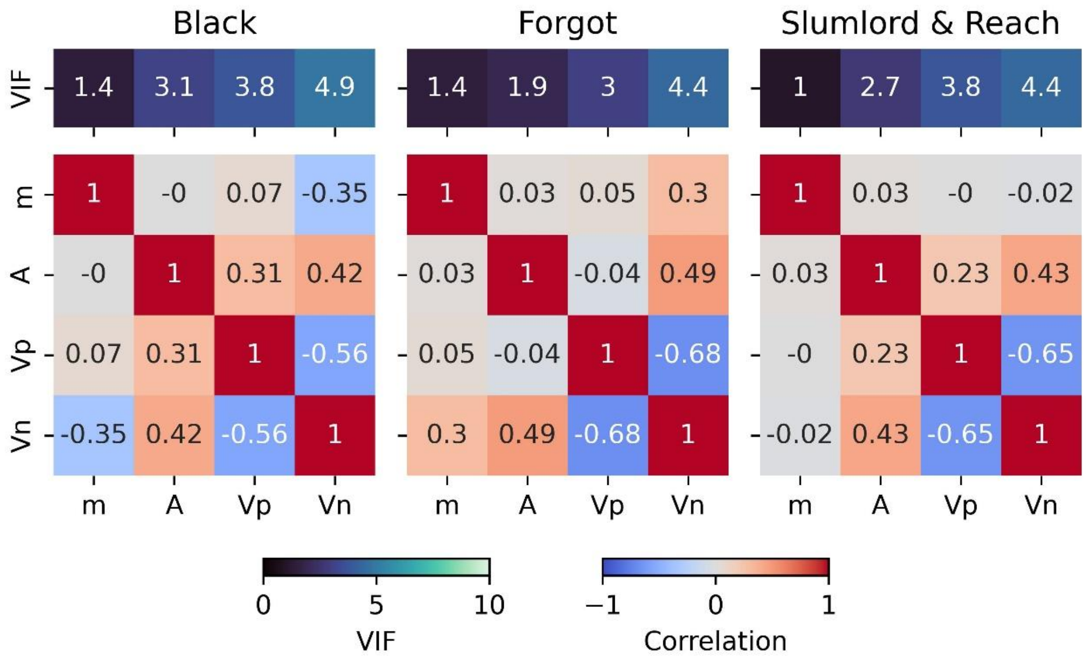


**Figure S5. Variance inflation factors (top) and inter-regressor correlations (bottom) for the four-parameter Bivalence model across stories.** *Notes.* m = the onset time regressor; A = the arousal regressor; Vp = the positive valence regressor; Vn = the negative valence regressor. Both valence and arousal ratings were mean-centered first. Valence ratings were computed using a Dual Rectified Rv-PCA procedure. All regressors were convolved with canonical HRF.

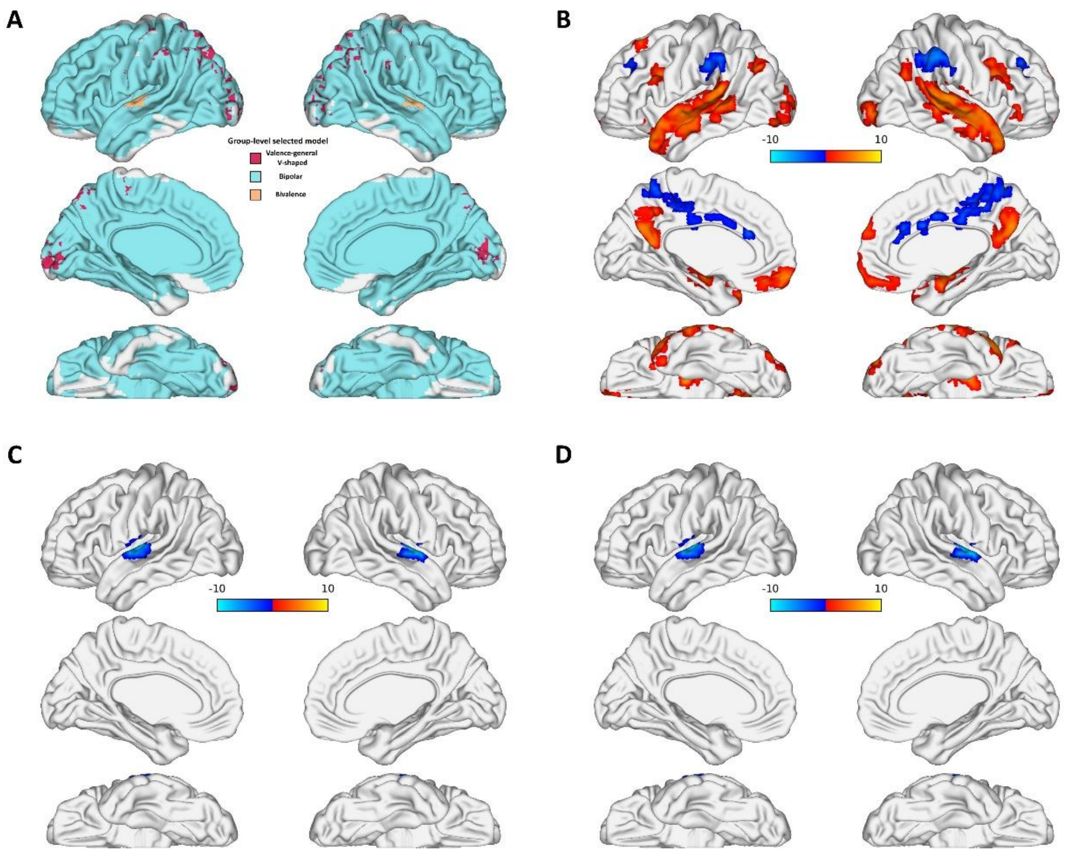


**Figure S6. Group-level cvBMS model selection and parametric modulation of valence in Model Family 3. A**. Group-level cvBMS map. **B**. Modulation effect of valence for the Bipolar model, after controlling for arousal. **C**. Modulation effect of positive valence for the Bivalence model, after controlling for arousal. **D**. Modulation effect of negative valence for the Bivalence model, after controlling for arousal. *Notes.* The results in panels B-D were shresholded using a cluster-size based correction, voxelwise $p < .001$.
